# A single-cell RNAseq atlas of the pathogenic stage of *Schistosoma mansoni* identifies a key regulator of blood feeding

**DOI:** 10.1101/2020.02.03.932004

**Authors:** George Wendt, Lu Zhao, Rui Chen, Chenxi Liu, Anthony J. O’Donoghue, Conor R. Caffrey, James J. Collins

## Abstract

Schistosomiasis is an ancient and chronic neglected tropical disease that infects over 240 million people and kills over 200,000 of the world’s poorest people every year^1, 2^. There are no vaccines and because there is only one drug available, the need for new therapeutics is great. The causative agents of this disease are flatworm parasites that dwell inside the host’s circulation, often for decades, where they feed on blood and lay eggs which are primarily responsible for disease pathology. As metazoans comprised of multiple tissue types, understanding the schistosome’s tissues on a molecular level and their functions during what can be decades of successful parasitism could suggest novel therapeutic strategies. Here, we employ single-cell RNAseq to characterize 43,642 cells from the pathogenic (adult) stage of the schistosome lifecycle. From these data, we characterize 68 molecularly distinct cell populations that comprise nearly all tissues described morphologically, including the nervous and reproductive systems. We further uncover a lineage of somatic stem cells responsible for producing and maintaining the parasite’s gut – the primary tissue responsible for digestion of host blood. Finally, we show that a homologue of *hepatocyte nuclear factor 4* (*hnf4*) is expressed in this gut lineage and required for gut maintenance, blood feeding and inducing egg-associated pathology *in vivo*. Together, the data highlight the utility of this single-cell RNAseq atlas to understand schistosome biology and identify potential therapeutic interventions.

---

Single-cell RNAseq (scRNAseq) is a powerful tool for comprehensively describing the various tissue types and basic physiology of diverse metazoans^3–6^. Although studies have used scRNAseq to describe non-pathogenic stages (*i.e.* larval and juvenile) of the schistosome lifecycle^7, 8^, the technology has not yet been employed to understand the biology of the pathogenic stage of schistosomes, or of any other metazoan parasite. To define the molecular signature of cell types in the adult schistosome, we dissociated adult *Schistosoma mansoni*, isolated cells by Fluorescence-Activated Cell Sorting (FACS), and generated scRNAseq libraries using a 10x genomics chromium controller (Fig. 1a). Schistosomes are unique among flatworms in that they are dioecious^9^ and sexual maturation of the female worm’s reproductive organs, including the ovary and vitellaria, requires close and sustained physical contact with the male worm^10^. Accordingly, to create a single cell atlas with the greatest diversity of cell types, we generated scRNAseq libraries from adult male parasites, adult sexually-mature female parasites, and age-matched virgin female parasites. Using these data, we performed unbiased clustering and identified 68 molecularly distinct clusters composed of 43,642 cells (Fig. 1b, Extended Data Fig. 1, Supplementary Table 1). These clusters included: three transcriptionally distinct clusters of proliferative cells that express the somatic stem cell (*i.e.,* neoblast) marker *nanos2*^11^ (Fig. 1c, Extended Data Fig. 2a); eight clusters of cells expressing markers of progenitor cells involved in generation of the schistosome tegument (“skin”-like surface)^12, 13^ (Extended Data Fig. 2b); two clusters of parenchymal cells (Fig. 1d, Extended Data Fig. 2c); one cluster of cells corresponding to ciliated flame cells that are part of the worm’s protonephridial (excretory) system (Fig. 1e, Extended Data Fig. 2d); eight separate clusters of muscle cells (Fig. 1f); and one cluster of oesophageal gland cells (Fig. 1g, Extended Data Fig. 2e). In spite of the theoretical difficulty in sorting syncytial cells by FACS, our analysis identified clusters of cells corresponding to known syncytial tissues, including the tegument^13, 14^ (Extended Fig. 2f) and gut^15^ (Fig. 1h, Extended Data Fig. 2g). However, we failed to identify cells from two other syncytial tissues, *i.e*., the female ootype (an organ involved in egg shell formation) and the protonephridial ducts (which are thought to be syncytial in other parasitic flatworms^16^) that together with the flame cells make up the protonephridial system^17^.

**Fig. 1.**
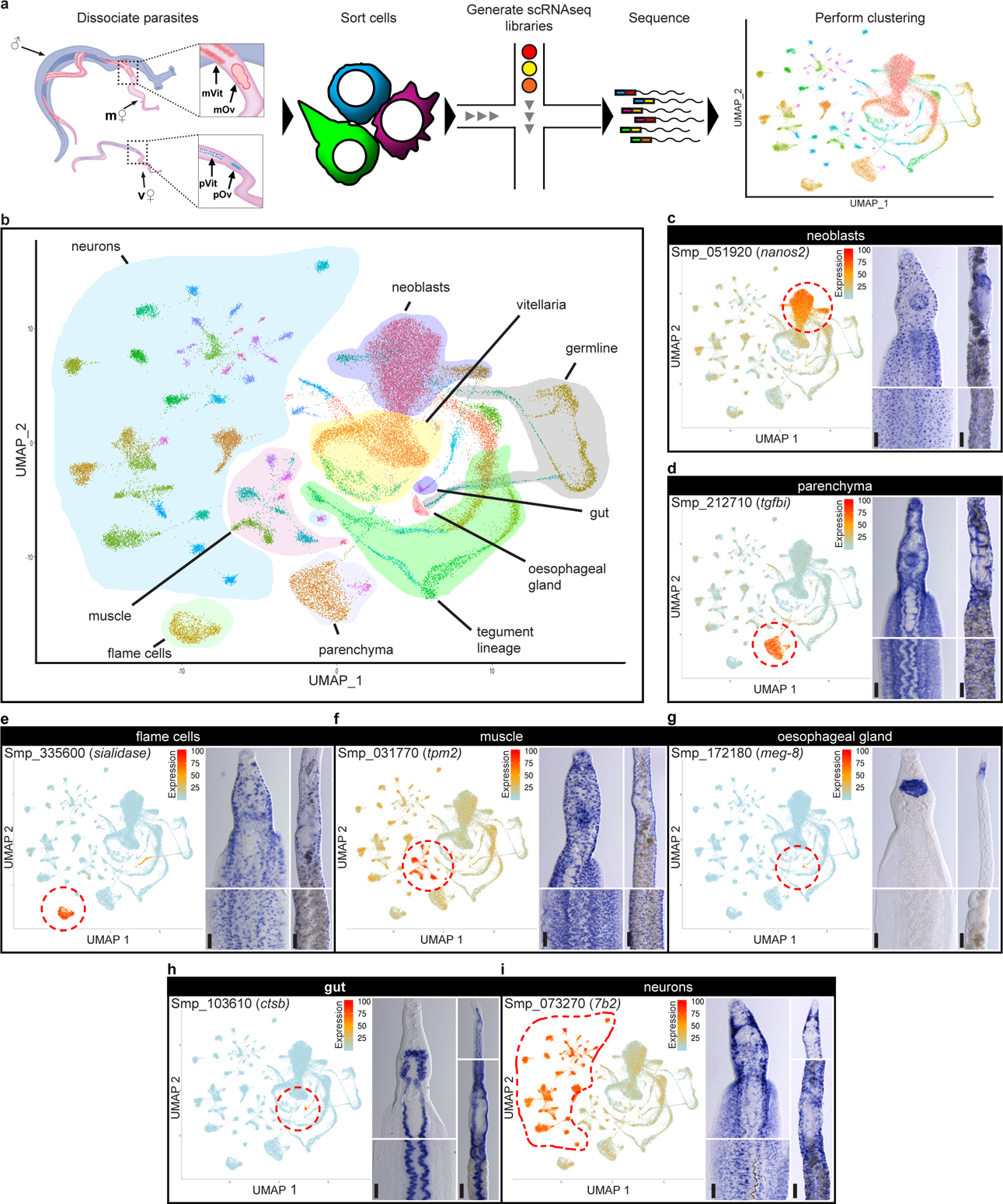
Overview of single-cell RNA sequencing of adult schistosome cells. a, Schematic diagram of single-cell RNA sequencing workflow. Cartoon to left depicts male paired with a mature female worm (m♀) that possess a mature ovary (mOv) and vitellaria (mVit); unpaired virgin female worms (v♀) possess a primordial ovary (pOv) and vitellaria (pVit). **b,** UMAP projection plot of the 68 clusters generated from the scRNAseq data. **c-i,** (top) UMAP projection plot and representative micrograph of colorimetric WISH of the indicated gene in the head (middle left) and body (middle right) of a male parasite and the ovary (bottom left) and vitellaria (bottom right) of a mature female parasite for the **c**, neoblast-specific gene *nanos2*, **d**, the parenchyma-specific gene *tgfbi,* **e**, the flame cell-specific gene *sialidase*, **f**, the muscle cell-specific gene *tpm2*, **g**, the oesophageal gland-specific gene *meg-8*, **h**, the gut-specific gene *ctsb*, and **i**, the neuron-specific gene *7b2*. Scale bars, all 100µm. UMAP projection plots colored by gene expression (blue = low, red = high).

**Fig. 2.**
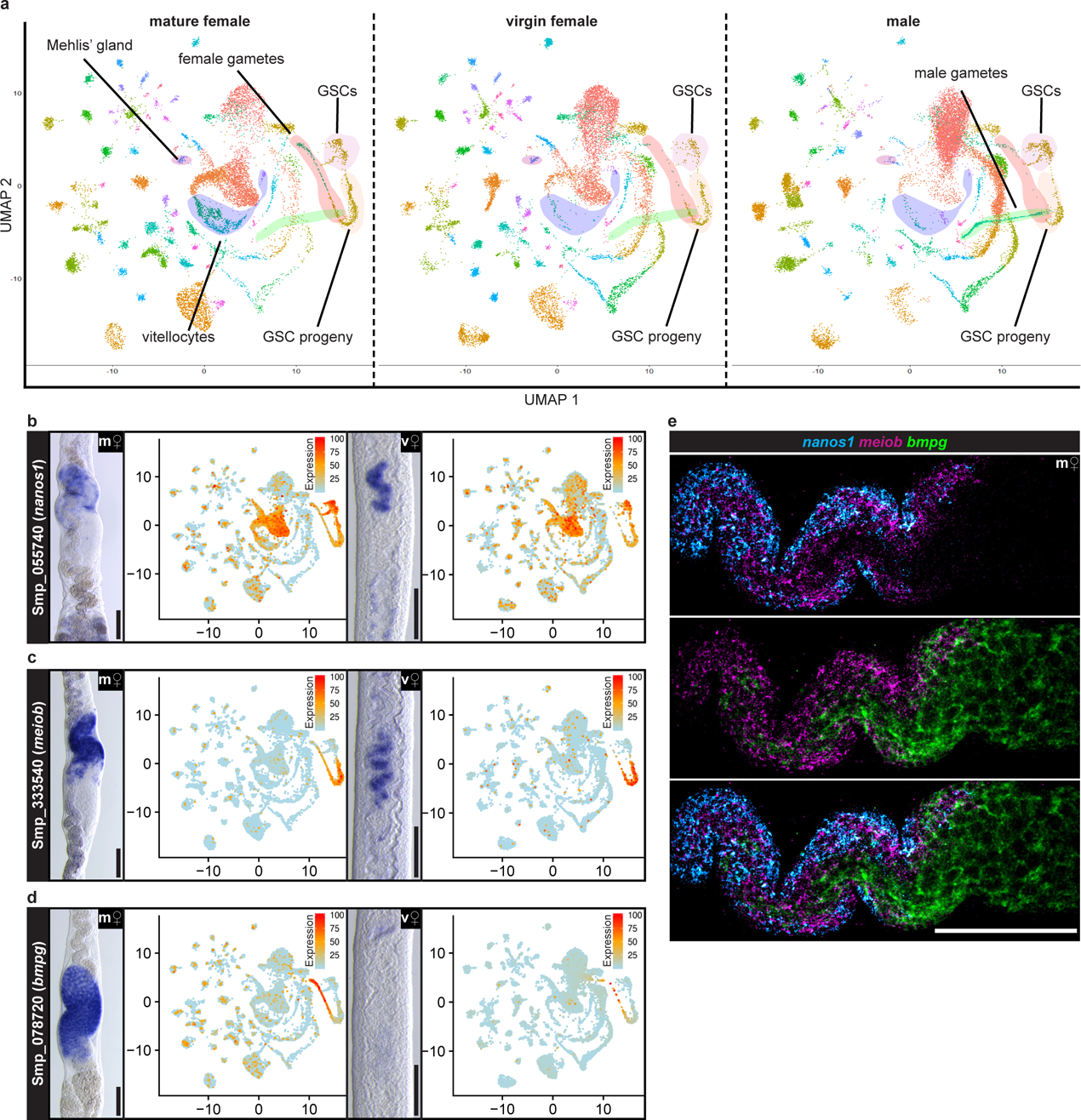
Identification of the germ lineage in schistosome ovary. a, UMAP projection plots of all clusters split by parasite sex. Sex-specific clusters are labeled. **b-d**, For the “GSCs”-enriched gene *nanos1* (**b**), the “GSC progeny”-enriched gene *meiob* (**c**), and the “female gametes”-enriched gene *bmpg* (**d**): (far left) representative micrograph of colorimetric WISH of indicated gene in sexually mature females (m♀), (mid left) UMAP projection plot of indicated gene expression in sexually mature females, (mid right) representative micrograph of colorimetric WISH of indicated gene in sexually immature females (v♀), and (far right) UMAP projection plot of indicated gene expression in sexually immature females. **e,** Representative micrograph of triple FISH of *nanos1*, *meiob* and *bmpg* in the ovary of a sexually mature female (m♀). Scale bars, all 100µm. UMAP projection plots are colored by gene expression (blue = low, red = high).

We uncovered a surprising level of molecular complexity within the schistosome nervous system, identifying 30 clusters of cells that express the neuroendocrine protein *7b2* (Fig. 1i, Extended Data Fig. 3a) and one apparent neuronal cluster of cells that did not express high-levels of *7b2* but expressed a variety of synaptic molecules, suggesting a neuron-like identity (Extended Data Fig. 3a, far right, Supplementary Table 1). Examination of genes from these neuronal cell clusters uncovered not only a highly-specific molecular fingerprint for several cell populations (Extended Data Fig. 3b, Supplementary Table 1), but evidence of a highly ordered structural and regional specialization in both the central and peripheral nervous systems, including evidence of left-right asymmetrical expression of a neuron-specific marker (Extended Data Fig. 3c) and as many as nine types of apparently ciliated neurons (Extended Data Fig. 3d,e). This complexity is rather surprising given the relatively “sedentary” lifestyle of adult parasites in the portal vasculature^9^. Further investigation of these various subpopulations of neurons could lead to discovery of novel mechanisms by which schistosomes perceive and interact with their environment.

**Fig. 3.**
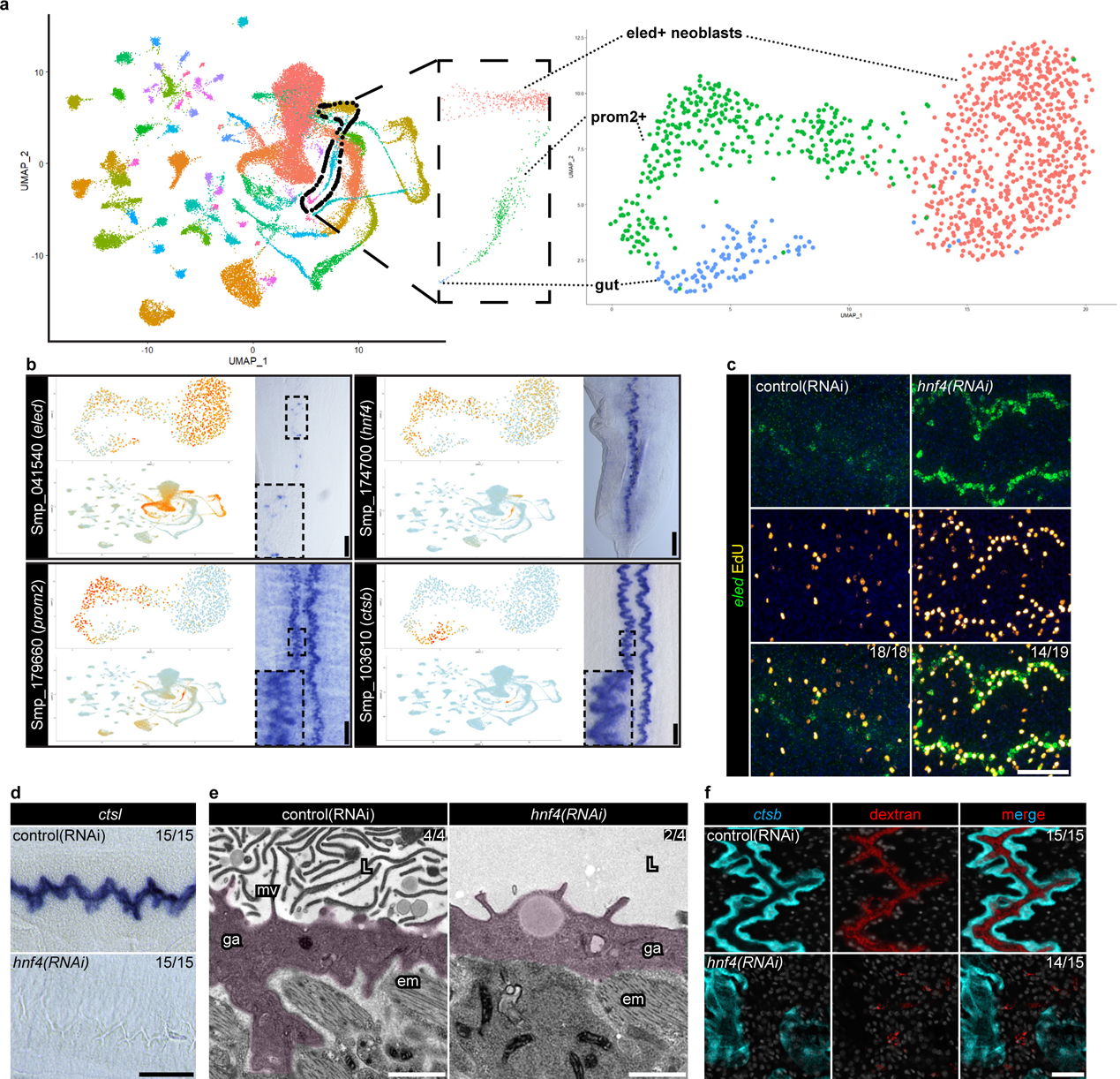
An *hnf4* homolog regulates a novel gut lineage **a**, Schematic of the re-clustering of the putative gut lineage from the single cell RNAseq data. **b**, (top) UMAP projection plots of the expression pattern of the indicated gene on the re-clustered dataset from Fig. 3a, (bottom) on the entire dataset, and (right) a representative micrograph from a colorimetric WISH of the parasite’s body for the putative gut neoblast marker *eled*, the putative gut progenitor marker *prom2*, the definitive gut marker *ctsb,* and the candidate gut-neoblast regulator *hnf4*. Insets show magnifications of the dashed boxes. **c**, Representative micrographs from FISH in conjunction with an EdU pulse showing the expression of *eled* (green) and presence of EdU^+^ proliferative cells (yellow) in either control RNAi conditions or *hnf4* RNAi conditions. The number of parasites similar to the representative micrograph is indicated in the upper right of the bottom panels. Data are from two biological replicates. Nuclei are pseudo-colored blue. **d**, Representative micrographs of colorimetric WISH of the “gut”-specific gene *ctsl* in either control RNAi conditions or *hnf4* RNAi conditions. The number of parasites grossly similar to the representative micrograph is indicated in the upper-right of each panel. Data are from three biological replicates. **e**, Representative TEM micrographs of the gut of either control(RNAi) or *hnf4(RNAi)* animals. The number of parasites similar to the representative micrograph is indicated in the upper right of each panel. Data are from four parasites from two biological replicates. ‘mv’ microvilli, ‘ga’ gastrodermis, ‘L’ lumen, ‘em’ enteric muscle. **f**, Representative micrographs from FISH showing the expression of *ctsb* (cyan) and the presence of fluorescently-labeled dextran (red) in the gut lumen in either control(RNAi) or *hnf4(RNAi)* animals. The number of parasites similar to the representative micrograph is indicated in the upper right of the far-right panels. Data are from three biological replicates. Nuclei are pseudo-colored grey. Scale bar, **b** 100µm, **c** 50µm, **d** 100µm, **e** 1µm, **f** 20µm. UMAP projection plots are colored by gene expression (blue = low, red = high).

Schistosome muscle is also very heterogeneous, with eight different clusters of cells that possess unique expression patterns (Extended Data Fig. 3f,g). Some populations appear to be diffusely arranged throughout the animal (“muscle 1” and “muscle 2”), whereas others are anatomically restricted such as the “muscle 7” cells that reside at the midline proximal to the parasite’s digestive tract, suggesting that this cluster represents cells of the enteric musculature (Extended Data Fig. 3f, third column).

Similar to what has been observed in planarians^18^, tapeworms^19^, and acoels^20^, we find that many well-characterized morphogens that regulate *wnt* (Extended Data Fig 4a-d) and *tgfb* signaling (Extended Data Fig. 4e-h) are predominantly expressed in muscle and neuronal cells. Homologues of many of these genes are expressed specifically in planarian muscles^3^ and have been implicated in regulating normal regeneration in planarians^18^. Though schistosomes survive amputation^21^, there is no evidence of whole-body tissue regeneration. It is interesting, therefore, that the expression pattern of these signaling molecules is conserved in a non-regenerative animal. This suggests that their anatomically restricted expression in neuromuscular tissues could regulate schistosome neoblast fates during homeostasis. Further investigation of this hypothesis in schistosomes could uncover novel regulators of stem cell biology in these parasites.

**Fig. 4.**
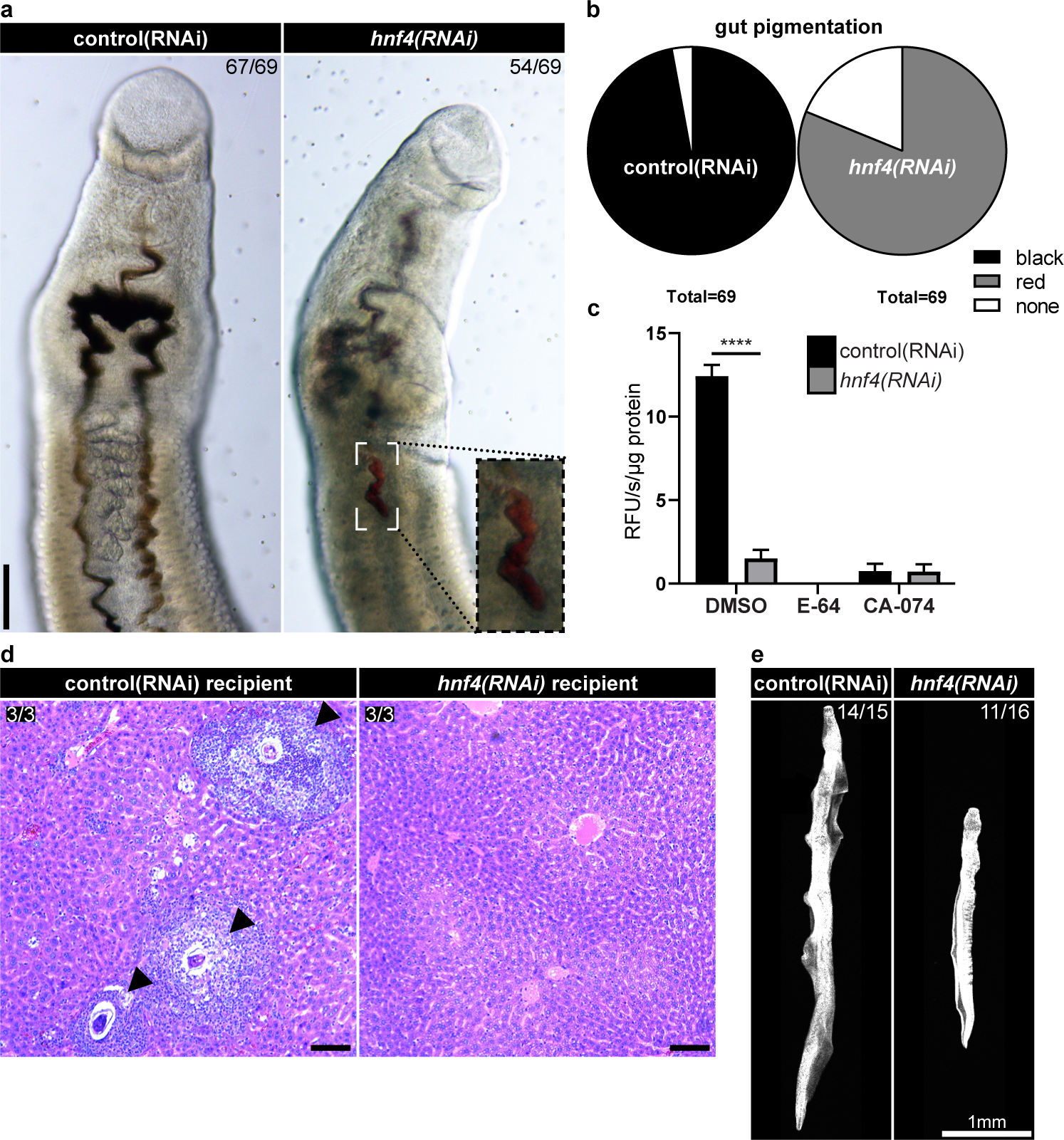
*hnf4* is required for blood feeding **a**, Representative brightfield micrographs of live control(RNAi) animals or *hnf4(RNAi)* animals that were cultured in media containing bovine red blood cells. The inset in the right panel shows a magnification of the indicated area of the gut filled with undigested hemoglobin as evidenced by the red pigmentation. The number of parasites similar to the representative micrograph is indicated in the upper right of each panel. **b**, Pie chart depicting the frequency of different gut pigmentation of animals from **a**. *n* = 69 control(RNAi) animals and 69 *hnf4(RNAi)* animals from three biological replicates. **c**, Graph of the cysteine protease cathepsin activity of lysates from control(RNAi) animals or *hnf4(RNAi)* animals as determined by the ability to cleave the fluorogenic substrate, Z-FR-AMC, in the presence of no inhibitor (DMSO), the general cysteine protease inhibitor, E-64, or the cathepsin B-selective inhibitor, CA-074. Data are from three biological replicates each in triplicate. **d**, Representative micrographs of H&E-stained sections of mouse livers 22 days after transplant with RNAi-treated parasites. No granulomata are present in the livers of mice that received *hnf4(RNAi)* parasites. The number of sections similar to the representative micrograph is indicated in the upper right each panel. Data are from three recipients from one biological replicate. **e**, Representative image of DAPI-stained parasites recovered from mice 22 days after transplant with RNAi-treated parasites. The number of parasites grossly similar to the representative micrograph is indicated in the upper right of each panel. Data are from parasites perfused from three separate recipients. Nuclei are pseudo-colored grey. Scale bars, **a**, 100µm, **e**, 100µm, **f**, 1mm. ****, *p*<0.0001 (Welch’s t-test).

The pathology of schistosome infection is driven almost exclusively by the host’s inflammatory responses to parasite eggs^22^. Therefore, understanding the biology of schistosome reproductive organs could lead to novel methods to target disease pathology. Our single-cell expression atlas allows us to study the differences between not only male and female parasites, but also between sexually mature and age-matched virgin females at the cellular level (Fig. 2a). Male, mature female, and age-matched virgin female parasites all have germline stem cells (GSCs) marked by expression of *nanos1*^23^. Our scRNAseq data revealed that GSCs have very similar gene expression patterns regardless of sex or maturity (Fig. 2b, Extended Data Fig. 5a). Much like GSCs, GSC progeny fall into the same clusters in both male and female parasites, suggesting no major sex- or maturation-dependent differences in early gametogenesis (Fig. 2c and Extended Data Fig 5b). However, mature gametes cluster according to sex, with substantial expression of “female gametes”-enriched genes only found in mature females (Fig. 2d and Extended Data Fig. 5c) and substantial expression of “male gametes”-enriched genes only found in males (Extended Data Fig. 5d).

Our scRNAseq data also enables us to study sexual cellular lineages. The sexually mature schistosome ovary is structured such that GSCs reside at the anterior pole whereas mature differentiated oocytes are found at the posterior end^23, 24^. The “GSCs”-enriched genes such as *nanos1* are expressed in the anterior ovary (Fig. 2b, left panel, Extended Data Fig. 5a, middle panels) and “female gametes”-enriched genes such as *bmpg* are expressed in the posterior ovary (Fig. 2d, left panel, Extended Data Fig. 5c, middle panels). Our single-cell RNAseq data shows that the “GSC progeny” cluster exists between “GSCs” and “female gametes” on the UMAP projection plot, (Fig. 2a), so we would expect the “GSC progeny”-enriched genes such as *meiob* to be expressed between the anterior and posterior ovary, which is indeed what we find (Fig. 2c, left panel, Extended Data Fig. 5b, middle panels). We would also predict proliferative cells to be concentrated in the *nanos1*^+^ GSCs, with little to no cell proliferation in *meiob*^+^ GSC progeny cells or *bmpg*^+^ female gametes, which agrees with our observations (Extended Data Fig. 6a-d). Concurrent visualization of ovarian stem cells, progenitors and oocytes reveals a highly-organized linear architecture (Fig. 2e). Interestingly, both mature and virgin females express the “GSC progeny” marker *meiob* (Fig. 2c), suggesting that the primordial ovary of the virgin female still undergoes some level of differentiation without stimulus from the male. Thus, it appears that male parasites may promote survival of differentiating GSCs rather than inducing GSC commitment. This observation is consistent with studies suggesting that male-female pairing can suppress apoptosis in the vitellaria of virgin female worms^25^. Further investigation to clarify exactly what happens to GSCs upon male pairing is required.

We were also able to use our single cell atlas to examine the schistosome vitellaria, another male-sensitive, stem-cell dependent tissue responsible for producing the yolk cells that provide nutrients to the parasite’s eggs. Despite a wholly different function and organization, there were many parallels between the maturation of the ovary and the vitellaria such as the presence of an apparent lineage from stem cell to mature tissue (Extended Data Fig. 6e-h). Our atlas also confirmed the decades-old observation that male parasites have a low frequency of vitellocyte-like cells^26^ (Extended Data Fig. 6e, bottom two panels). Finally, we identified markers of pairing-independent sexual tissues such as the flatworm-specific Mehlis’ gland that plays an enigmatic role in egg production^9^ (Extended Data Fig. 6i).

Previous work suggests that adult schistosome neoblasts are homogeneous and predominantly give rise to cells involved in tegument production^12, 13^. We identified a putative non-tegument lineage as suggested by a linear “path” of cells leading from a neoblast sub-population to the gut (Fig. 3a). The putative lineage began with a rare population of proliferative cells that expressed the somatic neoblast marker *nanos2* (Fig. 1c), the juvenile neoblast marker *eled*^7^ (Fig. 3b, top left, Extended Data Fig. 7a) and an *hnf4* homolog (Fig. 3b, top right, Extended Data Fig. 7b), but did not express the germ cell marker *nanos1* (Extended Data Fig. 7c). Adjacent to these *eled*^+^ neoblasts on the UMAP projection plot was the “*prom2*+” population, characterized by expression of *prom2* and *hnf4* in and around the gut (Fig. 3b, bottom left, Extended Data Fig 7d). Situated next to the “*prom2*+” cluster was the “gut” cluster, which expressed definitive gut markers such as genes encoding cathepsin B-like cysteine proteases *ctsb* (Fig. 3b, bottom right, Extended Data Fig. 7e).

Based on the localization of these genes on the UMAP projection plot (Fig. 3a), their expression patterns, and that *hnf4* is a marker of gut stem cells in planarians^27^, we hypothesized that the *eled*^+^ neoblasts, “*prom2+*” cells, and “gut” cells represent the schistosome gut lineage. In order to test this model, we sought to perturb the *eled*^+^ neoblasts at the top of lineage in order to observe the effects on downstream cells. To this end, we performed a small-scale RNAi screen targeting several genes expressed in the *eled*^+^ neoblasts (Extended Data Fig. 8a, b). Remarkably, RNAi of *hnf4* resulted in massive expansion of *eled*^+^ neoblasts along the parasite’s gut (∼3.8-fold increase in *hnf4(RNAi)* animals compared to control, p < 0.0001) (Fig. 3c, Extended Data Fig. 8c-f). According to our lineage model, an expansion of *eled*^+^ neoblasts could either result in an increase in gut production because of an expanded stem cell pool, or it could result in a decrease in gut production because of a differentiation block. Supporting the second model, we found expression of several definitive gut markers such as *ctsl* (Smp_343260) and *ctsb* (Smp_103610) were decreased upon *hnf4* RNAi (Fig. 3d, Extended Data Fig. 9a). We next performed *in situ* hybridization (ISH) to examine the localization of transcripts and found that several gut transcripts were no longer expressed, their expression was reduced, or their expression pattern was significantly altered (Extended Data Fig. 9b). To examine the extent of gut dysfunction, we performed RNAseq on *hnf4(RNAi)* animals. We found that over 70% of all transcripts expressed in the “gut” cluster in our single-cell data set were significantly downregulated following *hnf4* RNAi (Extended Data Fig. 9c, Supplementary Table 2). Indeed, a look at the top 25 most downregulated genes in the RNAseq experiment revealed that all were expressed in the gut and 21 were expressed almost exclusively in the gut (Extended Data Fig. 9d).

To determine whether these transcriptional changes in *hnf4(RNAi)* animals affected the gut structure, we examined *hnf4(RNAi)* animals by transmission electron microscopy (TEM). The schistosome gut is a syncytial blind tube-like structure with a microvilli-filled lumen^15^. Though gut tissue was still present, we found a significant decrease in luminal microvilli (Fig. 3e, Extended Data Fig. 9e). Additionally, we found that 2 out of 4 of *hnf4(RNAi)* animals had massively dilated lumens compared to 0 out of 4 of control(RNAi) animals (Extended Data Fig. 9f, f’). To understand whether the parasites were capable of filling their gut lumen, we supplemented the culture media with fluorescently-labeled dextran that, upon ingestion, enters the gut lumen, and is absorbed into the gut in a time-dependent fashion^28^. After 12 hours of culture in dextran (after dextran is ingested but before it is absorbed), 15 out of 15 control parasites had dextran in the gut lumen whereas only 1 out of 15 *hnf4(RNAi)* parasites had dextran in the lumen (Fig. 3f). Further examination of the parasite’s head revealed that dextran completely failed to enter the digestive tract of the *hnf4(RNAi)* parasites (Extended Data Fig. 9g), suggesting either a complete loss of patency or a defect in the parasite’s ability to coordinate the passage of dextran into the gut.

Although the gut was abnormal in *hnf4(RNAi)* animals, it was unclear whether the *hnf4* RNAi resulted in destruction of the gut, a block in new gut production or some combination of both. There was no apparent difference in the number of TUNEL^+^ apoptotic cells between control and *hnf4(RNAi)* animals (Extended Data Fig. 9h). To understand whether stem cell differentiation was grossly intact, we looked at tegument production using EdU pulse-chase approaches in *hnf4(RNAi)* animals and found a significant increase in tegument production compared to control(RNAi) animals (Extended Data Fig. 9i, j), ruling out a broad stem cell differentiation defect. Our ability to monitor new gut production by EdU pulse-chase approaches was complicated by the fact that gut marker expression was largely absent in most parasites (Fig. 3d, Extended Data Fig. 9b). Examination of gut differentiation in cases where we could detect gut marker expression by EdU pulse-chase approaches in *hnf4(RNAi)* parasites revealed that new gut-like tissue (*i.e*., expresses gut markers like *ctsb*, though not always in the typical linear pattern along the parasite’s midline) was still being produced (Extended Data Fig. 9k), but the gut-like tissue that was present was morphologically abnormal by ISH (Extended Data Fig. 9b, see *hnf4(RNAi)* animals). Examination of *eled* expression in conjunction with the gut marker *ctsb* revealed that areas with a greater number of *eled^+^* cells had low or no expression of *ctsb*. Conversely, where *ctsb* transcripts were present, *eled*^+^ cells were relatively sparse (Extended Data 9l). This is consistent with a partial differentiation block where areas with fewer *eled^+^* cells and high *ctsb* expression may represent locations where *eled*^+^ neoblasts were able to partially overcome the differentiation block and form gut-like tissue. However, given the relatively low basal rate of gut production^12^, a partial block of gut differentiation is not likely to result in such a dramatic gut defect over the course of a 17 day RNAi treatment. As such, *hnf4* is likely required for both normal gut production and maintenance.

Based on the profound morphological defects in the gut, we next asked whether there were any functional consequences of *hnf4* RNAi. Although glucose can be absorbed across the parasite’s tegument, parasites rely on the gut to digest host blood cells^29^. To test the digestive capability of *hnf4(RNAi)* parasites, we added red blood cells to the media and observed the parasites’ ability to uptake and digest the cells. While the vast majority of control(RNAi) parasites (67/69) were able to ingest and digest red blood cells as evidenced by black pigmentation in the gut^30^, *hnf4(RNAi)* parasites either failed to ingest red blood cells (15/69) or ingested red blood cells but couldn’t digest them as evidenced by red pigmentation in the gut (54/69) (Fig. 4a, b). These data suggest a decrease in the blood ingestion and digestion capacity of the *hnf4(RNAi)* animals but does not address the mechanism of any digestive defects. Because we measured a decrease in the expression of many proteolytic enzymes in our RNAseq experiment (Supplementary Table 2), we next asked whether there was a loss in the *hnf4(RNAi)* parasites of those cysteine (cathepsin) proteases that contribute to hemoglobin digestion^31^. Accordingly, we measured the cathepsin activity of lysates from control(RNAi) and *hnf4(RNAi)* parasites using the fluorogenic peptidyl substrate, z-Phe-Arg-AMC (Z-FR-AMC)^32^. We show that the majority of the activity (94%) in protein extracts from control(RNAi) parasites is due to cathepsin B, as this activity is sensitive to the selective cathepsin B inhibitor, CA-074 (Fig 4c). In *hnf4(RNAi)* parasites, the cysteine protease activity is decreased 8.2-fold relative to control(RNAi) parasites. Thus, the functional assay data are consistent with our gene expression analyses that show a significant reduction in five cathepsin B gene sequences in the *hnf4*(RNAi) animals (Supplementary Table 2). In contrast, we show that aspartyl protease activity is unchanged in control(RNAi) and *hnf4(RNAi)* parasites (Extended Data Fig. 10a), which could reflect the expression of aspartic proteases in non-gut tissues that were not downregulated following *hnf4* RNAi (Supplementary Table 1, 2). Taken together, these data suggest that *hnf4* is required for cathepsin B-mediated digestion of hemoglobin in *S. mansoni*.

Given the importance of blood uptake and digestion for egg production^29^, the primary driver of the pathology of schistosomiasis, we wondered whether *hnf4* was required to cause disease in the host. To test this, we transplanted control(RNAi) and *hnf4(RNAi)* parasites into uninfected mice and then perfused the mice 23 to 30 days post-transplant. Worm recovery was statistically indistinguishable (72% of control(RNAi) animals recovered vs. 49% of *hnf4(RNAi)* animals, *p* = 0.136) (Extended Data Fig. 10b), suggesting no substantial impact on parasite survival. Nonetheless, mice transplanted with *hnf4(RNAi)* parasites had grossly normal looking livers in contrast to abundant egg-induced granulomata in control(RNAi) recipients (Fig. 4d, Extended Data Fig. 10c). Also, recovered male *hnf4(RNAi)* parasites were significantly shorter than their control counterparts (2.87mm vs. 5.21mm, respectively, *p* < 0.0001) (Fig. 4e, Extended Data Fig. 10d). Together, these results suggest that *hnf4* is required for parasite growth and egg-induced pathology *in vivo*.

Schistosomiasis is a neglected tropical disease due in no small part to the difficulty of studying these parasites in the laboratory. Prior to this work, identification of specific tissue markers and understanding the cellular and molecular consequences of experimental perturbations relied upon a great deal of effort and guesswork^7, 12, 13, 17, 23, 33^. Using scRNAseq, we not only generated the most comprehensive single-cell atlas of any metazoan parasite to date, but also identified regulators of gut biology, leveraging this knowledge to experimentally perturb schistosome-induced pathology in the mammalian host. Indeed, our approach serves as a template for the investigation of other understudied and experimentally challenging parasitic metazoans, thereby improving our understanding of their biology and enabling us to discover novel therapies for these pathogens.

## Materials and Methods Parasite acquisition and culture

Adult *S*. *mansoni* (NMRI strain, 6–7 weeks post-infection) were obtained from infected female mice by hepatic portal vein perfusion with 37°C DMEM (Sigma-Aldrich, St. Louis, MO) plus 10% Serum (either Fetal Calf Serum or Horse Serum) and heparin. Parasites were cultured as previously described ^12^. Unless otherwise noted, all experiments were performed with male parasites. Experiments with and care of vertebrate animals were performed in accordance with protocols approved by the Institutional Animal Care and Use Committee (IACUC) of UT Southwestern Medical Center (approval APN: 2017-102092).

## Fluorescence Activated Cell Sorting

FACS sorting was performed as previously described^13^ with minor modifications. Freshly perfused adult male and sexually mature adult female worms were separated by incubation in a 0.25% solution of tricaine ^11^ for approximately 5 minutes. Sexually immature adult virgin female worms were separately perfused from single-sex infected mice. Male, mature female, or virgin female worms were suspended in a 0.5% solution of Trypsin/EDTA (Sigma T4174) in PBS. The worms were then triturated for approximately 15 minutes until the solution became turbid and no large pieces of worms were left. The trypsin was inactivated by adding an equal volume of serum-containing media. The dissociated worms were then centrifuged at 500 *g* for 10 m at 4°C. Next the worms were resuspended in 1 ml of Basch media with 10 µL of RQ1 DNAse (Promega M6101) and incubated for 10 minutes at RT. The dissociated worms were centrifuged again at 500 *g* for 10 minutes at 4°C. The cells were then resuspended in 1mL of staining media (0.2% BSA, 2mM EDTA in PBS, pH7.40) and incubated in Hoechst 33342 (18 µg/ml) (Sigma B2261) for 1 hour at RT in the dark. 9mL of staining media was then added to the worms and then the whole suspension was filtered through a 40 µm cell strainer. The worms were centrifuged once again at 500 *g* for 10 minutes at 4°C. Worms were then resuspended in 1mL of staining media containing Hoechst 33342 (18 µg/ml) and propidium iodide (1 µg/ml) (Sigma-Aldrich P4170) and then filtered once more through a 40 µm cell strainer into a 12×75mm FACS tube. Filtered cells were then sorted on a FACSAria II custom (BD Biosystems) with 305/405/488/561/633nm lasers. Sorts were performed with a 100 µm nozzle and cells were sorted into sorting media (0.2% BSA in PBS, pH7.40). For all FACS experiments, a Hoechst threshold was applied to exclude debris and improve the efficiency of sorting.

## Single-cell RNA sequencing

FACS-sorted cells were centrifuged again at 500 *g* for 10 minutes at 4°C then resuspended in 0.2% BSA in PBS. Libraries were created using a Chromium Controller (10x Genomics) according to manufacturer guidelines and sequenced in using a NextSeq 500 (illumina). Sequencing data was processed and mapped to the *Schistosoma mansoni* genome (v7) using Cell Ranger (10x Genomics). Unfiltered data from Cell Ranger was imported into Seurat (v3.1.1)^34, 35^ and cells were filtered as follows: Female (nFeature_RNA (> 750), nCount_RNA (1500-20000), Percent Mitochondrial (<3%); Male/Virgin female (nFeature_RNA (> 750), nCount_RNA (1000-20000), Percent Mitochondrial (<3%)). Mitochondrial genes were identified as those with the prefix “Smp_9”. Each of the 9 individual datasets (Supplementary Table 3) was normalized (NormalizeData) and variable features were identified (FindVariableFeatures, selection.method = “vst”, nfeatures = 2000). From here, integration anchors were identified (FindIntegrationAnchors, dims 1:78), the data was integrated (IntegrateData, dims = 1:78, features.to.integrate = features), and scaled (ScaleData). We then ran RunPCA, RunUMAP (reduction = “pca”, dims = 1:78, n.neighbors = 40), FindNeighbors (reduction = “pca”, dims = 1:78), FindClusters (resolution = 5). The number of principal components (78) used for this analysis was defined by JackStraw. Analysis of the resulting single cell map found that clusters 27 and 50 contained few enriched markers, therefore we removed the 964 cells present in these clusters and reran the analysis with 78 principal components. From here we generated the final UMAP projection plot with RunUMAP (n.neighbors = 36, min.dist = 0.70, dims = 1:80). Next, we generated clusters (FindClusters, resolution = 5) and manually inspected the unique genes expressed in each of the clusters. In some cases we found that some of the 85 resulting clusters did not express a core set of unique genes, therefore, these clusters were merged into a single cluster of cells as follows: Neoblasts (clusters 0,1,2,6,7,37), Neoblast progeny (cluster 4,8), Neuron 1 (clusters 10, 60, 68), Neuron 6 (clusters 24, 26), Parenchyma (clusters 11, 12, 51), flame cells (clusters 14, 41), S1 Cells (clusters 3, 9, 32, 42) and tegument (clusters 36, 63). After merging we were left with a final map of 68 clusters of 43,643 cells. Raw data from single cell RNAseq experiments are available from XXXX with accession number XXXX.

## Parasite labeling and imaging

Colorimetric and fluorescence in situ hybridization analyses were performed as previously described ^11, 12^ with the following modification. To improve signal-to-noise for colorimetric in situ hybridization, all probes were used at 10 ng/mL in hybridization buffer*. In vitro* EdU labeling and detection was performed as previously described^11^. For dextran labeling of the parasite gut, 10 male RNAi-treated parasites were given 10µL/mL of 5 mg/mL (in water) solution of biotin-TAMRA-dextran (Life Technologies D3312) and cultured 12 hours. The parasites were then fixed in fixative solution (4% formaldehyde in PBSTx (PBS + 0.3% triton-X100)) for 4 hours in the dark with mild agitation. Worms were then washed with 10 ml of fresh PBSTx for 10 minutes, then dehydrated in 100% methanol and stored at −20dC until used in fluorescence in situ hybridization as described^11, 12^. All fluorescently labeled parasites were counterstained with DAPI (1 µg/ml), cleared in 80% glycerol, and mounted on slides with Vectashield (Vector Laboratories).

Transmission electron microscopy samples were prepared from RNAi-treated parasites that were immersed in fixative (2.5% glutaraldehyde in 0.1M sodium cacodylate buffer pH 7.4 with 2mM CaCl2) and then amputated at the head and the tail in order to retain ∼5mm of trunk. After three rinses with 0.1 M sodium cacodylate buffer, the parasite trunks were embedded in 3% agarose and sliced into small blocks (1mm^3^), rinsed with the fixative three times and post-fixed with 1% osmium tetroxide and 0.8 % Potassium Ferricyanide in 0.1 M sodium cacodylate buffer for one and a half hours at room temperature. Samples were rinsed with water and *en bloc* stained with 4% uranyl acetate in 50% ethanol for two hours. They were then dehydrated with increasing concentration of ethanol, transitioned into propylene oxide, infiltrated with Embed-812 resin and polymerized in a 60°C oven overnight. Blocks were sectioned with a diamond knife (Diatome) on a Leica Ultracut 7 ultramicrotome (Leica Microsystems) and collected onto copper grids, post stained with 2% aqueous Uranyl acetate and lead citrate. Images were acquired on a Tecnai G2 spirit transmission electron microscope (FEI, Hillsboro, OR) equipped with a LaB6 source at 120kV using a Gatan ultrascan CCD camera.

Blood in the parasite gut and Haematoxylin and Eosin stained-samples were imaged with brightfield light using a Zeiss AxioZoom V16 equipped with a transmitted light base and a Zeiss AxioCam 105 Color camera.

Confocal imaging of fluorescently labeled samples was performed on a Nikon A1 Laser Scanning Confocal Microscope. Unless otherwise mentioned, all fluorescence images represent maximum intensity projection plots. To perform cell counts, cells were manually counted in maximum intensity projection plots derived from confocal stacks. In order to normalize counts, we collected confocal stacks and normalized the number of cells counted to the length of the parasite in the imaged region. Brightfield images were acquired on a Zeiss AxioZoom V16 equipped with a transmitted light base and a Zeiss AxioCam 105 Color camera.

## RNA interference

For detailed schematic of RNAi experiments, see Supplementary Table 4. Generally, all experiments utilized freshly perfused male parasites (separated from females) unless otherwise noted. dsRNA treatments were all carried out at 30µg/ml in Basch Media 169. dsRNA was generated by *in vitro* transcription and was replaced as indicated in Supplementary Table 4. EdU pulses were performed at 5µM for 4 hours before either fixation or chase as previously described^11^.

As a negative control for RNAi experiments, we used a non-specific dsRNA containing two bacterial genes ^36^. cDNAs used for RNAi and in situ hybridization analyses were cloned as previously described ^36^; oligonucleotide primer sequences are listed in Supplementary Table 5.

## qPCR and RNAseq

RNA collection was performed as previously described^12^ with the following modifications. Parasites were treated with dsRNA as described in Supplementary Table 3 (“strategy 4”) and whole parasites were collected in Trizol. RNA was purified from samples utilizing Direct-zol RNA miniprep kits (Zymo Research R2051). Quantitative PCR analyses were performed as previously described ^11, 12^. cDNA was synthesized using iScript™ cDNA synthesis kit (Bio-Rad 1708891) and qPCR was performed as previously described^13^ utilizing iTaq™ Universal SYBR® Green Supermix (Bio-Rad 1725122) and a QuantStudio 3 Real-Time PCR System (Applied Biosystems); oligonucleotide primer sequences used for qPCR are listed in Supplementary Table 5. RNAseq on *hnf4(RNAi)* parasites was performed as previously described^13^ using TruSeq Stranded mRNA Library Prep (illumina 20020594) to prepare libraries, which were sequenced on a NextSeq 550 (illumina). The total number of reads per gene was determined by mapping the reads to the *<u>S. mansoni</u>* genome (v7) using STAR (version 020201)^37^. S. mansoni genome sequence and GTF files used for mapping were acquired from Wormbase Parasite^38^. Pairwise comparisons of differential gene expression were performed with DESeq2 (version 1.12.2)^39^. Volcano plots were made with using the “volc” function from ggplot2. In order to filter out noise, genes with a base-mean expression value less than 50 were excluded from analysis. Furthermore, genes that were differentially expressed (padj < 0.05) that were not assigned to the automatically assigned to the “gut” cluster during initial clustering were manually examined in the single-cell RNAseq data and those that were expressed in the gut were reclassified to the “gut” cluster. Raw data from *hnf4* RNAi RNAseq experiments are available at XXXX with accession number XXXX.

## Protease activity assays

To measure cysteine protease cathepsin activity^32^, five worms of each RNAi condition (see Supplementary Table 3 “strategy 7”) were ground and sonicated in 300 µL assay buffer (0.1 M citrate-phosphate, pH 5.5). The lysate was centrifuged at 15,000*g* for 5 minutes and the pellet was discarded. The total protein concentration was calculated using the bicinchoninic acid assay with bovine serum albumin as the protein standard. Each well in the assay had 1 µg of protein. The assay buffer was 0.1 M citrate-phosphate, pH 5.5 with 2 mM DTT. CA-074 (Cayman Chemical, 24679-500) and E-64 (Alfa Aesar, J62933) controls were set up by incubating the sample with 10 µM of each inhibitor for 30 min at room temperature. The final substrate concentration of Z-FR-AMC (R&D Systems, ES009) was 10 µM. The release of the AMC fluorophore was recorded in a Synergy HTX multi-mode reader (BioTek Instruments, Winooski, VT) with excitation and emission wavelengths at 340nm and 460nm, respectively.

To measure aspartic protease cathepsin activity, five worms of each RNAi condition (See Table 3 “strategy 7”) were ground and sonicated in 300 µL assay buffer (0.1 M citrate-phosphate, pH 5.5). The lysate was centrifuged at 15,000*g* for 5 mins and the pellet was discarded. Each well in the assay had 1 µg of protein. The assay buffer was 0.1 M citrate-phosphate, pH 3.5. Pepstatin A (MP Biomedicals, 0219536805) and E-64 controls were set up by incubating the sample with 10µM of either inhibitor for 30 minutes at room temperature. The final substrate concentration of mca-GKPILFFRL-K(dnp) (CPC Scientific, SUBS-017A) was 10µM. The release of the AMC fluorophore was recorded in a Synergy HTX multi-mode reader (BioTek Instruments, Winooski, VT) with excitation and emission wavelengths at 320nm and 400nm, respectively.

All protease activity experiments were carried out as biological triplicates each in triplicate.

## Surgical transplantation of schistosomes

Surgical transplantation was performed as previously described^40^ with the following modifications. Seven days prior to surgery, 5-week-old parasites were recovered from mice and treated with 30 µg/ml dsRNA for 7 days in Basch Media 1 69 (see Table 3 “strategy 8”). Before mice were anesthetized, 10 pairs (male and female) were sucked into a 1ml syringe, the syringe was fitted with a custom 25G extra thin wall hypodermic needle (Cadence, Cranston, RI), the air and all but ∼200 µL of media were purged from the needle, and the syringe was placed needle down in a test tube to settle the parasites to the bottom of the syringe. Mice were kept on infrared heating pads for the duration of the surgery. Following wound closure, mice received a single subcutaneous 20 µL dose of a 1 mg/mL solution of Buprenorphine SR-LAB CIII for analgesia and were allowed to recover on a warm heating pad. Mice were group housed and individual mice were tracked by ear punches. On either day 22 or day 30 post-transplantation mice were sacrificed and perfused to recover parasites. Male and female parasites were counted and fixed for 4 hours in 4% formaldehyde in PBSTx. Recipient livers were removed and fixed for 72 hours in 4% formaldehyde in PBS. The percentage parasite recovery was determined by dividing the total (male and female) number of worms transplanted by the total number of parasites recovered following perfusion. Livers from individual mice were sectioned and processed for Haematoxylin and Eosin staining by the UT Southwestern Molecular Pathology Core.

## Statistical analysis

All two-way comparisons were analyzed using Welch’s t-test. All three-way comparisons were analyzed using one-way ANOVA. RNAseq data was analyzed by the Wald test in DeSeq2. *p* values are indicated in the figure legends or in Supplementary Table 2.

## Acknowledgements

This work was supported by the National Institutes of Health R01 R01AI121037 (JJC), R21 R21AI133393 (AJOD), and F30 1F30AI131509-01A1 (GRW), the Welch Foundation I-1948-20180324 (JJC), the Burroughs Wellcome Fund (JJC), the Wellcome Trust 107475/Z/15/Z (JJC), and the Bill and Melinda Gates Foundation OPP1171488 (CRC). Some schistosome-infected mice were provided by the NIAID Schistosomiasis Resource Center for distribution through BEI Resources, NIH-NIAID Contract HHSN272201700014I. NIH: *Schistosoma mansoni*, Strain NMRI, Exposed Swiss Webster Mice, NR-21963. FACS was performed with the aid of the Moody Foundation Flow Cytometry Facility at the University of Texas Southwestern Medical Center (UTSW). TEM imaging and sample preparation was performed with the aid of the Electron Microscopy Core at UTSW. RNAseq was performed with the aid of the McDermott Center Next Generations Sequencing Core at UTSW.

---

**Extended Data Fig. 1.**
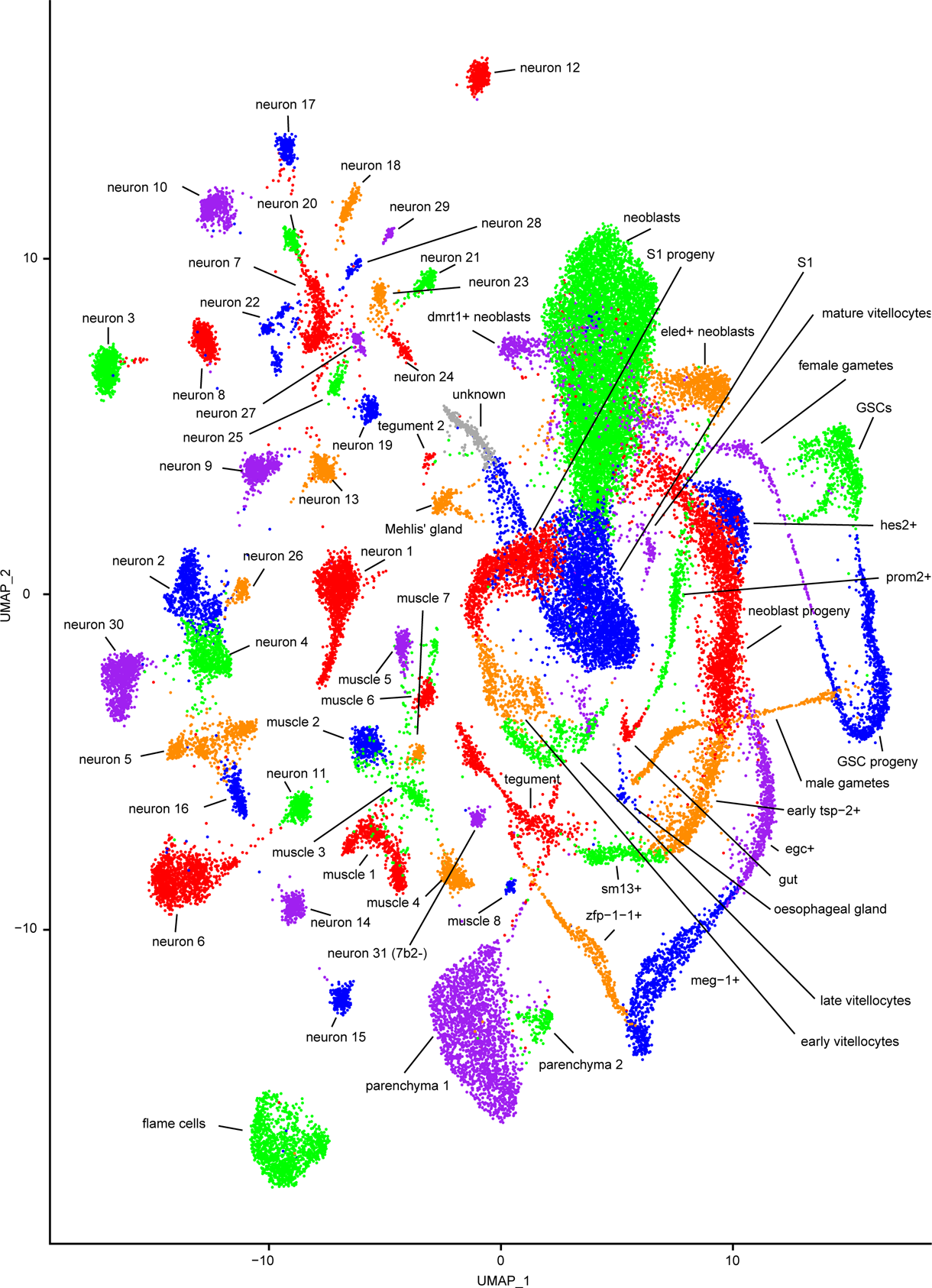
UMAP projection of all clusters with labels. Labeled UMAP projection plot of 68 clusters of cells from adult male, sexually mature adult female, and sexually immature adult virgin female *Schistosoma mansoni*.

**Extended Data Fig. 2.**
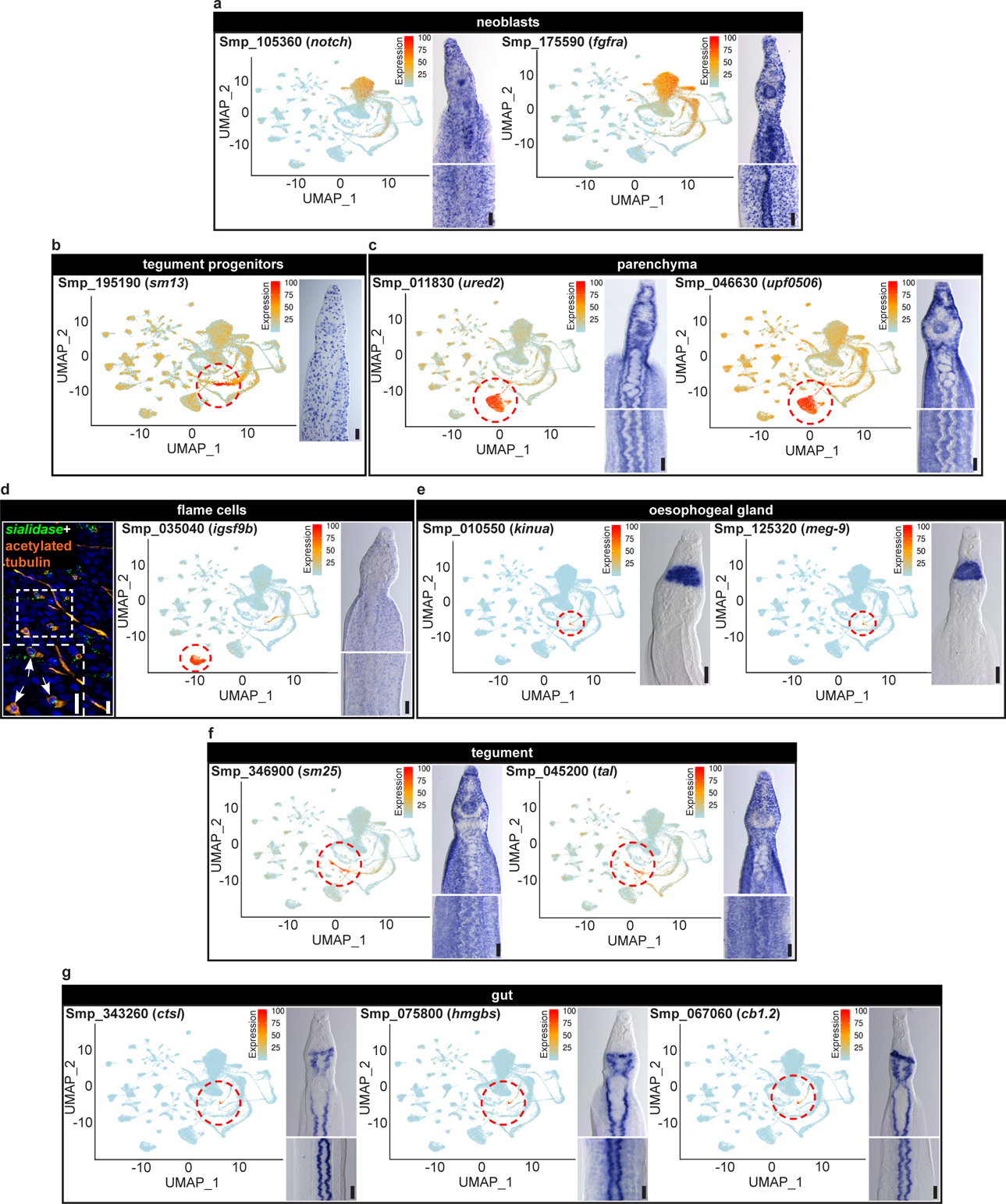
Additional somatic tissue-specific genes. a, (left) UMAP projection plot and (right) representative micrograph of colorimetric WISH of neoblast-specific genes *notch* and *fgfra*. **b,** (left) UMAP projection plot and (right) representative micrograph of colorimetric WISH of tegument progenitor-specific gene *sm13*. **c**, (left) UMAP projection plot and (right) representative micrograph of colorimetric WISH of parenchyma-specific genes *ured2* and *upf0506*. **d**, (left) representative micrograph of FISH in combination with acetylated tubulin immunofluorescence to label cilia, (middle) UMAP projection plot, and (right) representative micrograph of colorimetric WISH of flame cell-specific gene *igsf9b*. **e**, (left) UMAP projection plot and (right) representative micrograph of colorimetric WISH of oesophageal gland-specific genes *kinua* and *meg-9*. **f**, (left) UMAP projection plot and (right) representative micrograph of colorimetric WISH of tegument-specific genes *sm25* and *tal*. **g**, (left) UMAP projection plot and (right) representative micrograph of colorimetric WISH of gut-specific genes *ctsl, hmgbs,* and *cb1.2*. Scale bars, **d**, left panel: 10 µm. All others: 100µm. UMAP projection plots are colored by gene expression (blue = low, red = high).

**Extended Data Fig. 3.**
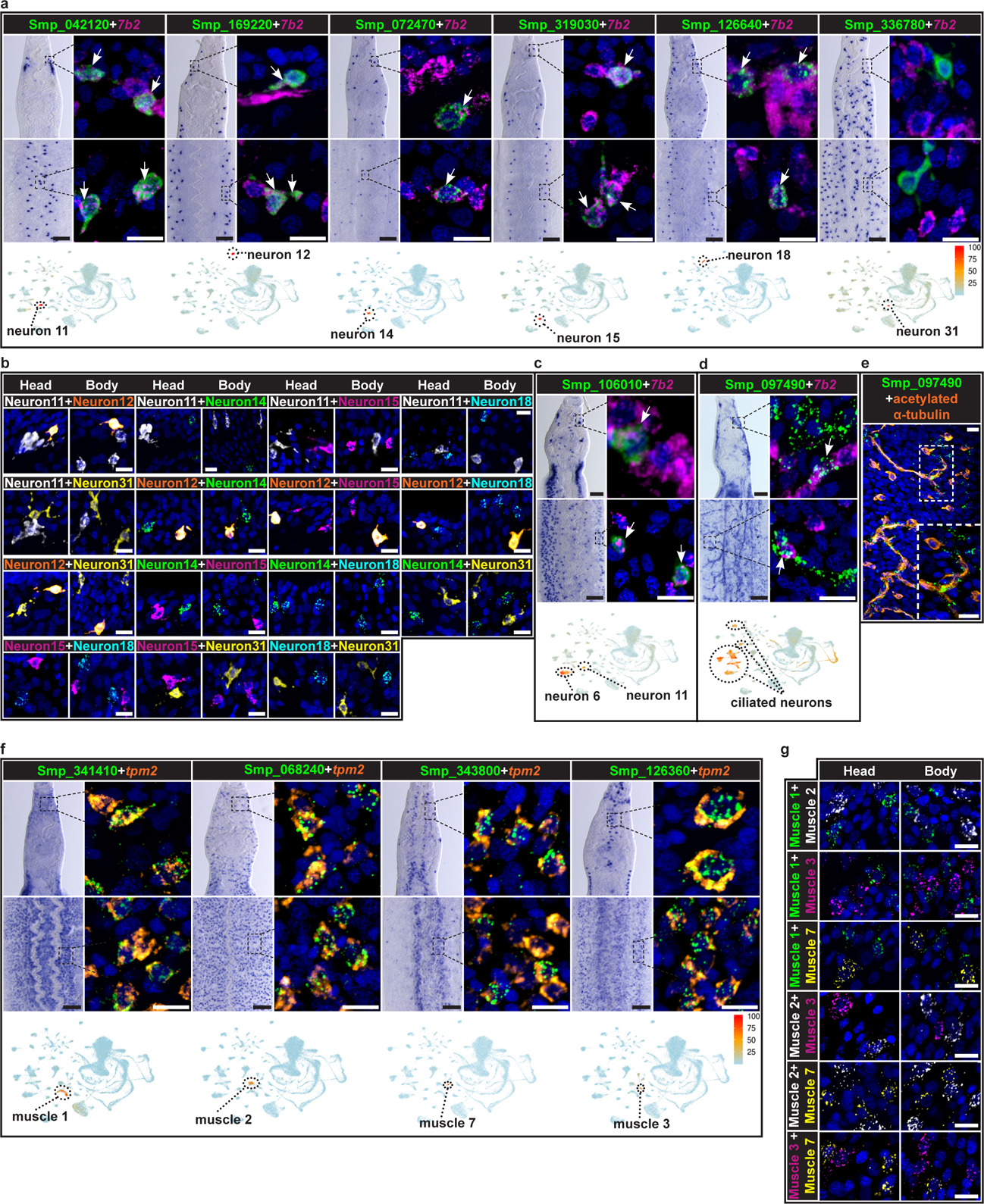
Schistosome muscles and neurons display complex heterogeneity. a, For each of 6 different neuron cluster-specific genes (from left to right “neuron 11”: *Smp_042120*, “neuron 12”: *Smp_159220*, “neuron 14”: *Smp_072470*, “neuron 15”: *Smp_319030*, “neuron 18”: *Smp_126640*, and “neuron 31”: *Smp_336780)*: (top left) representative micrograph of colorimetric WISH of head, (top right) representative micrograph of double FISH of region of head indicated in colorimetric WISH with cluster specific gene (green) and *7b2* (magenta), (middle left) representative micrograph of colorimetric WISH of body, (middle right) representative micrograph of double FISH of region of body indicated in colorimetric WISH with cluster specific gene (green) and *7b2* (magenta), and (bottom) UMAP projection plot. **b,** representative micrographs of double FISH with the indicated neuron cluster-specific markers showing no overlapping expression. **c-d,** (top left) representative micrograph of colorimetric WISH of head, (top right) representative micrograph of double FISH of region of head indicated in colorimetric WISH with **c**, “neuron 6”- and “neuron 11”-enriched gene *Smp_106010* (green) or **d**, the ciliated neuron-enriched gene *Smp_097490* (green) and *7b2* (magenta), (middle left) representative micrograph of colorimetric WISH of body, (middle right) representative micrograph of double FISH of region of body indicated in colorimetric WISH with *Smp_106010* (green) and *7b2* (magenta), and (bottom) UMAP projection plot. **e**, Representative micrograph of FISH of *Smp_097490* (green) with immunofluorescent labeling of acetylated tubulin (orange) and (bottom) UMAP projection plot. **f,** For each of 4 different muscle cluster-specific genes (from left to right “muscle 1”: *Smp_341410*, “muscle 2”: *Smp_068240*, “muscle 7”: *Smp_343800*, and “muscle 3”: *Smp_126360*: (top left) representative micrograph of colorimetric WISH of head, (top right) representative micrograph of double FISH of region of head indicated in colorimetric WISH with cluster specific gene (green) and the pan muscle marker *tropomyosin2* (*tpm2,* orange), (middle left) representative micrograph of colorimetric WISH of body, (middle right) representative micrograph of double FISH of region of body indicated in colorimetric WISH with cluster specific gene (green) and *tpm2* (orange), and (bottom) UMAP projection plot. **g**, Representative micrograph of double FISH with the indicated muscle cluster-specific genes showing no overlapping expression. Nuclei are pseudo-colored blue in all images. Scale bars, all FISH: 10µm, all colorimetric WISH: 100µm. UMAP projection plots are colored by gene expression (blue = low, red = high).

**Extended Data Fig. 4.**
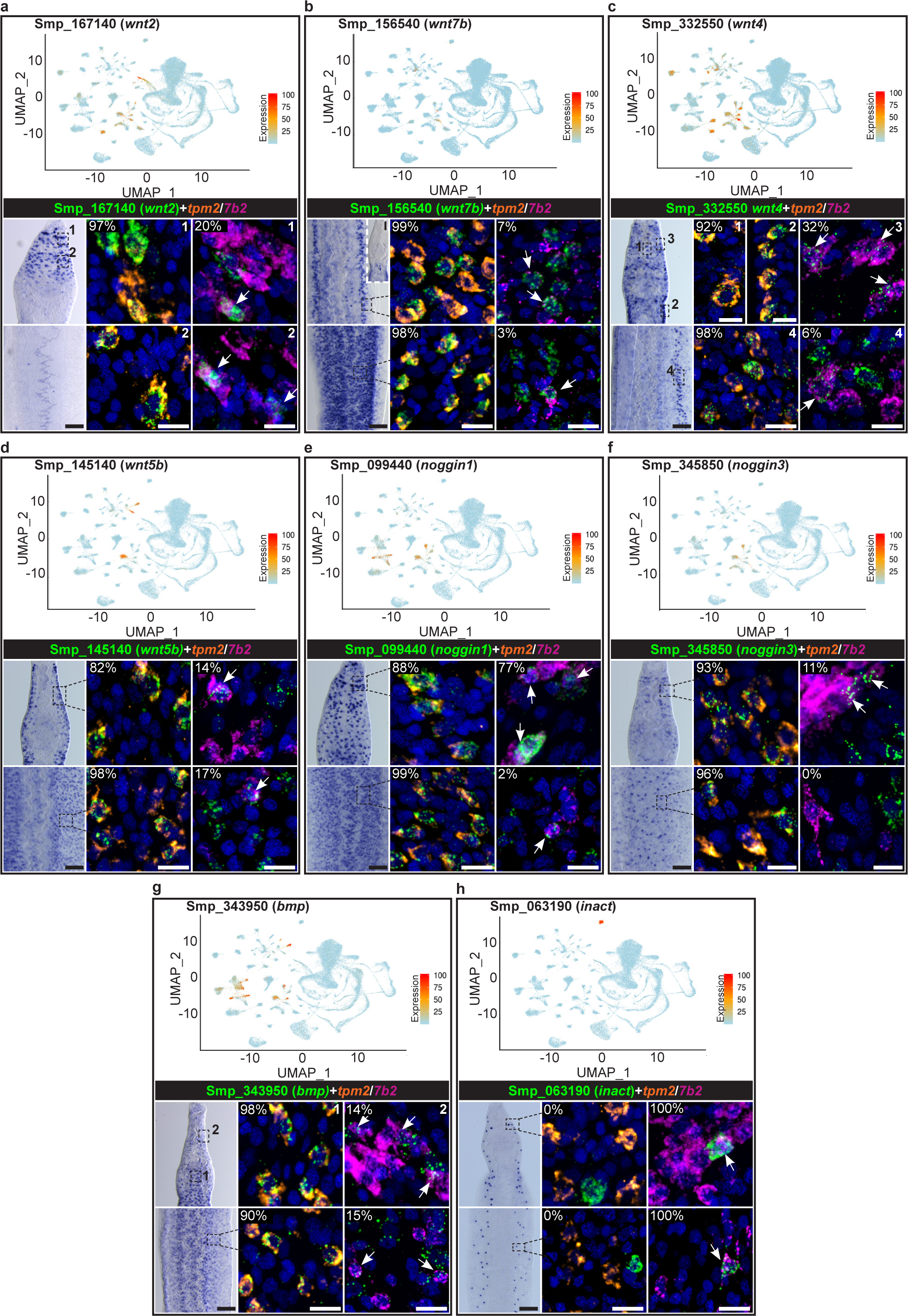
Morphogen homologs are expressed in schistosome muscles and neurons. For all genes: (top) UMAP projection plots, (middle left) representative micrograph of colorimetric WISH of head, (middle middle) representative micrograph of double FISH of region indicated in colorimetric WISH with muscle-specific gene *tpm2*, (middle right) representative micrograph of double FISH of region indicated in colorimetric WISH with neuron-specific gene *7b2*, (bottom left) representative micrograph of WISH of body, (bottom middle) representative micrograph of double FISH of region indicated in WISH with muscle-specific gene *tpm2*, (bottom right) representative micrograph of double FISH of region indicated in colorimetric WISH with neuron-specific gene *7b2* for wnt pathway genes **a**, *Smp_167140* (*wnt2*), **b**, *Smp_156540* (*wnt7b*), **c**, *Smp_332550* (*wnt4*), and **d**, *Smp_145140* (*wnt5b*) or tgfβ pathway genes **e**, *Smp_099440* (*noggin1*), **f**, *Smp_345850* (*noggin3*), **g**, *Smp_343950* (*bmp*), and **h**, *Smp_063190* (*inact*). Percentage in upper left corner of micrographs indicates percent of co-expression of the indicated gene with either *tpm2* or *7b2*. *n* = >100 cells from 3 different animals for all counts. Nuclei are pseudo-colored blue in all images. Scale bars, all FISH: 10µm. all WISH: 100µm. UMAP projection plots are colored by gene expression (blue = low, red = high).

**Extended Data Fig. 5.**
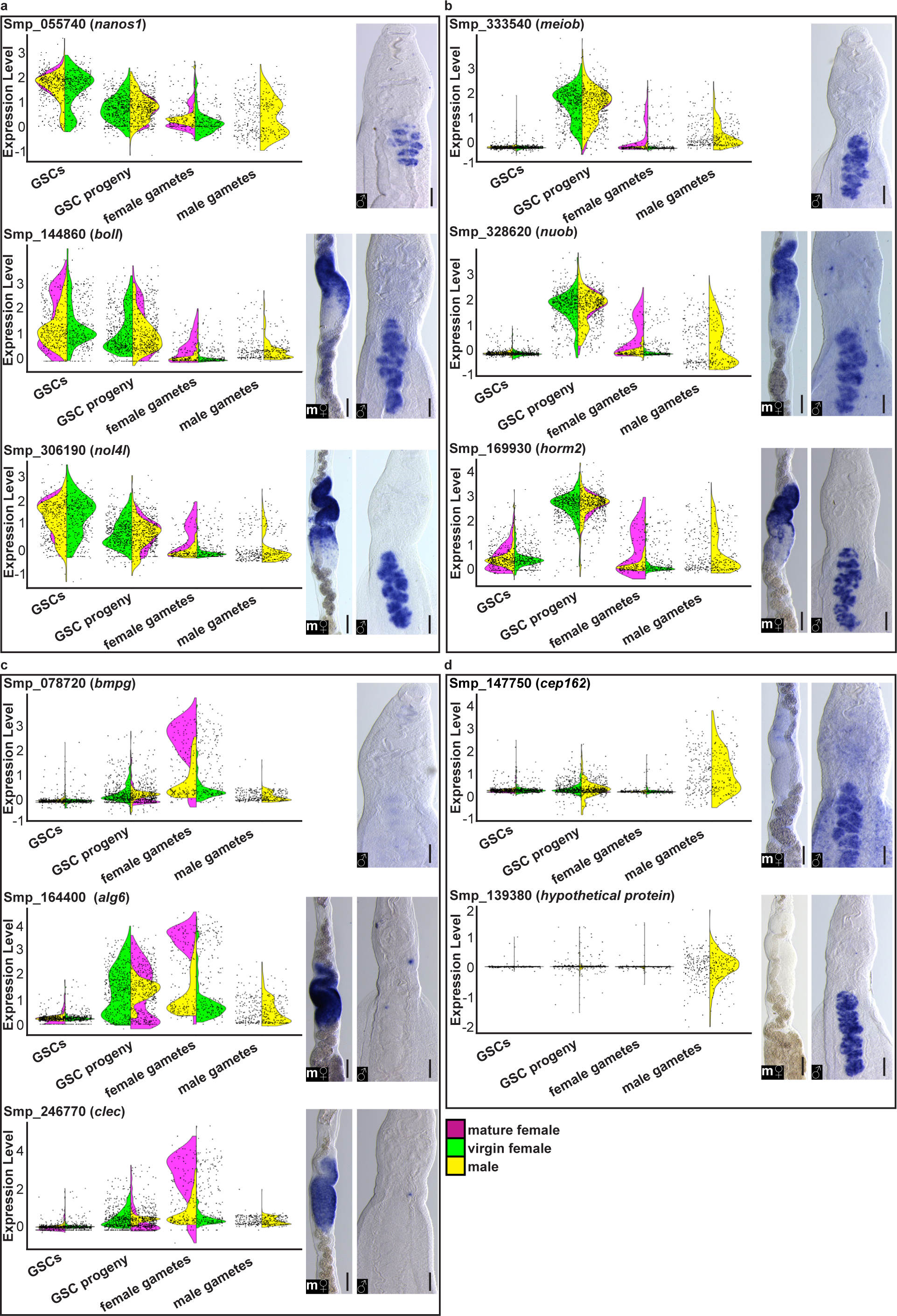
Additional reproductive tissue-specific genes. a-d, For the **a**, “GSCs”-enriched genes *nanos1*, *boll*, and *nol4l*, **b**, “GSC progeny”-enriched genes *meiob, nuob,* and *horm2*, **c**, “female gametes”-enriched genes *bmpg, alg6,* and *clec*, and **d**, “male gametes”-enriched genes *cep162* and *Smp_139380*: (left) violin plots showing gene expression levels across the clusters “GSCs”, “GSC progeny”, “female gametes”, “male gametes” colored by sex (mature female = magenta, virgin female = green, male = yellow) and (middle and right) representative micrographs of colorimetric WISH of the indicated gene in the (middle) ovary of sexually mature females (m♀) and (right) testes of males (♂). Scale bars, all 100µm.

**Extended Data Fig. 6.**
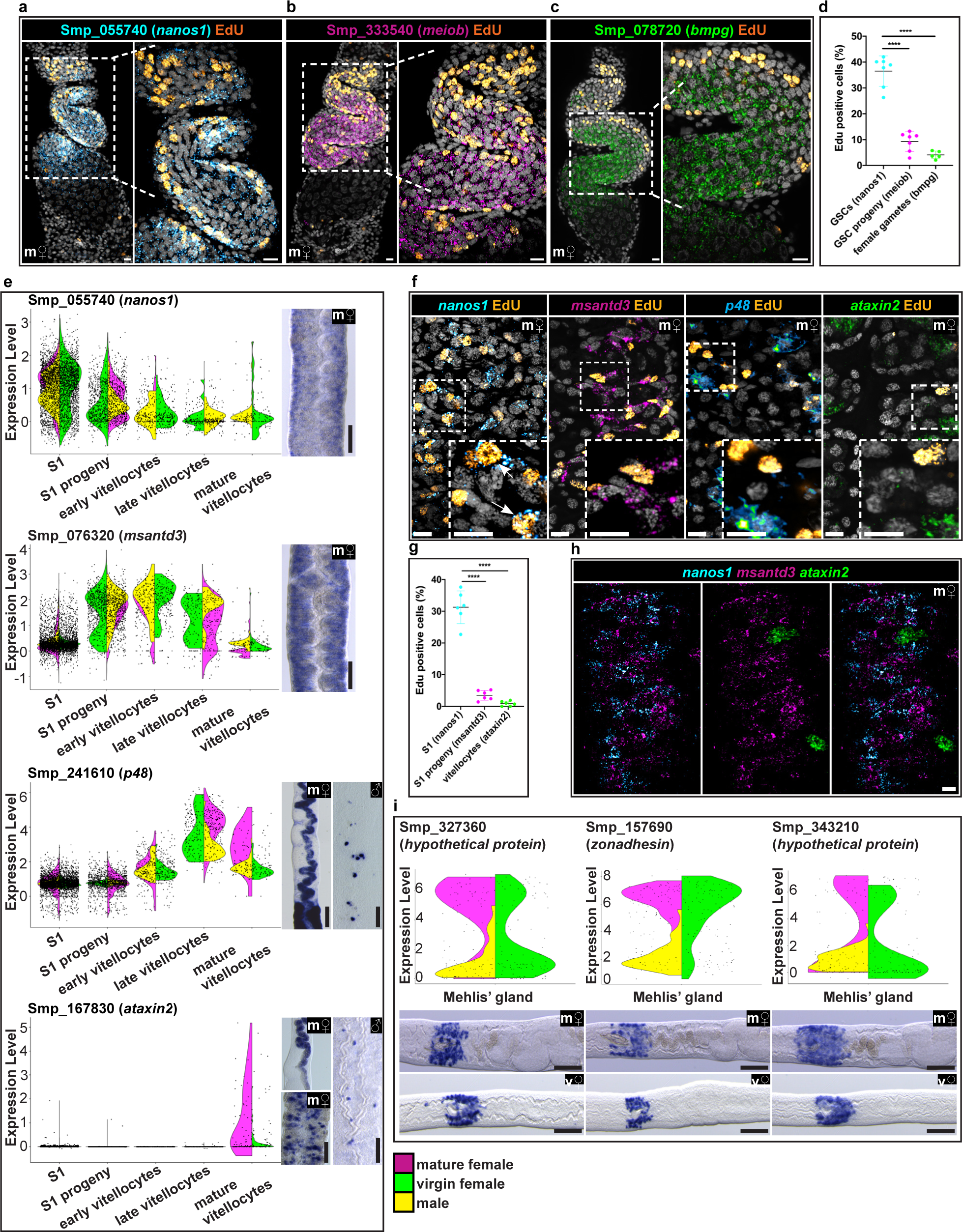
EdU labeling of proliferative cells in the ovary and vitellaria. a-c, Representative micrograph of FISH of the GSC marker *nanos1* (cyan) (**a**), the “GSC progeny”-enriched gene *meiob* (magenta) (**b**), or the “female gamete”-enriched gene *bmpg* (green) (**c**) in conjunction with a 30-minute EdU pulse (orange) to label the actively proliferating cells of the ovary of a sexually mature female (m♀). Nuclei are pseudo-colored grey. **d**, Graph showing quantification of percentage of *nanos1*^+^, *meiob*^+,^ or *bmpg*^+^ cells that are EdU^+^ following a 30-minute EdU pulse. **e**, For the “S1”-enriched gene *nanos1*, the “S1 progeny”-enriched gene *msantd3*, the “late vitellocyte”-enriched gene *p48*, and the “mature vitellocyte”-enriched gene *ataxin2*: (left) violin plots showing gene expression levels across the clusters “S1”, “S1 progeny”, “early vitellocytes”, “late vitellocytes”, “mature vitellocytes” colored by sex (mature female = magenta, virgin female = green, male = yellow) and (right) representative micrographs of colorimetric WISH of the indicated gene in the vitellaria of mature females (m♀) and the midline of males (♂) as indicated on the image. **f**, For the “S1”-enriched gene *nanos1*, the “S1 progeny”-enriched gene *msantd3*, the “late vitellocyte”-enriched gene *p48*, and the “mature vitellocyte”-enriched gene *ataxin2*: Representative micrograph of FISH for indicated gene (cyan/magenta/cyan-hot/green, respectively) in conjunction with an EdU pulse to show the localization of proliferative cells (orange) in the vitellaria of a sexually mature female. Nuclei are pseudo-colored grey. **g**, Graph showing quantification of percentage of *nanos1*^+^, *meiob*^+,^ or *bmpg*^+^ cells that are EdU^+^ following a 30-minute EdU pulse. *nanos1* percent EdU^+^. **h**, Representative micrograph of triple FISH of *nanos1*, *msantd3* and *ataxin2* in the vitellaria of a sexually mature female. **i**, For the “Mehlis’ gland”-enriched gene *Smp_327360*, *zonadhesin*, and *Smp_343210*: (top) violin plots showing gene expression levels in the “Mehlis gland” cluster colored by sex (mature female = magenta, virgin female = green, male = yellow) and (bottom) representative micrographs of colorimetric WISH of the indicated gene in region anterior to the ovary in sexually mature females (m♀) and virgin females (v♀) as indicated on the image. Scale bars, **a-c**, 10µm, **e**, 100µm, **f**, 10µm, **h**, 10µm, **i**, 100µm. ****, *p*<0.0001 (one-way ANOVA test).

**Extended Data Fig. 7.**
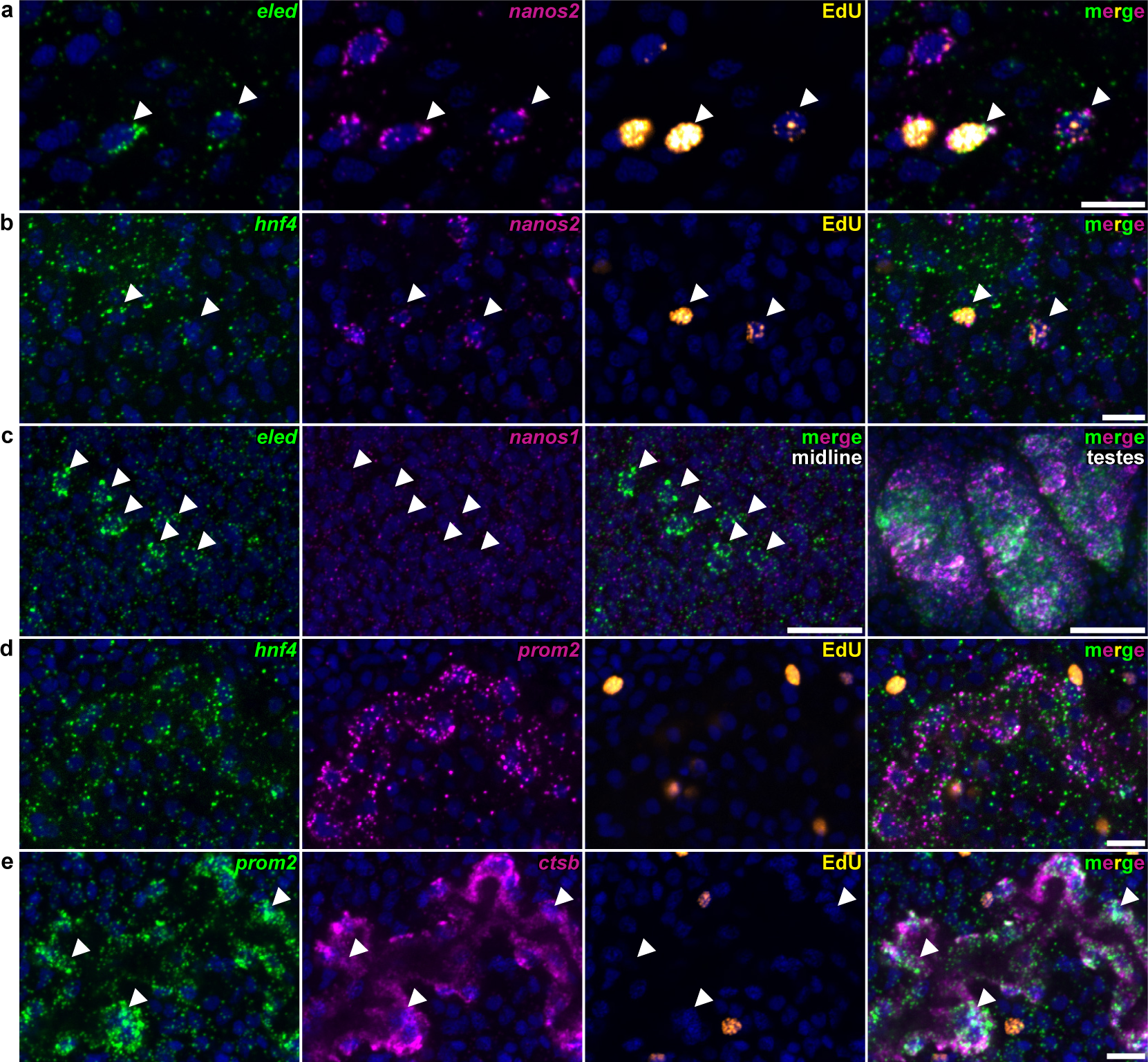
A putative schistosome gut lineage **a**, Representative micrographs of double FISH of *eled* and *nanos2* in EdU^+^ proliferative cells demonstrating co-expression. **b**, Representative micrographs of double FISH of *hnf4* and the pan-neoblast marker *nanos2* in EdU^+^ proliferative cells demonstrating co-expression. **c**, Representative micrographs of double FISH of *eled* and the GSC marker *nanos1* demonstrating no co-expression along the parasite’s midline but strong co-expression of *eled* and *nanos1* in the testes. **d**, Representative micrographs of double FISH of *hnf4* and *prom2* demonstrating co-expression in a gut-like pattern along the parasite’s midline. **e,** Representative micrographs of double FISH of *prom2* with the gut marker *ctsb* demonstrating the co-expression. Regions of high *prom2* expression with low *ctsb* expression are indicated with arrow heads. Nuclei are pseudo-colored blue in all images. Scale bars, 10µm.

**Extended Data Fig. 8.**
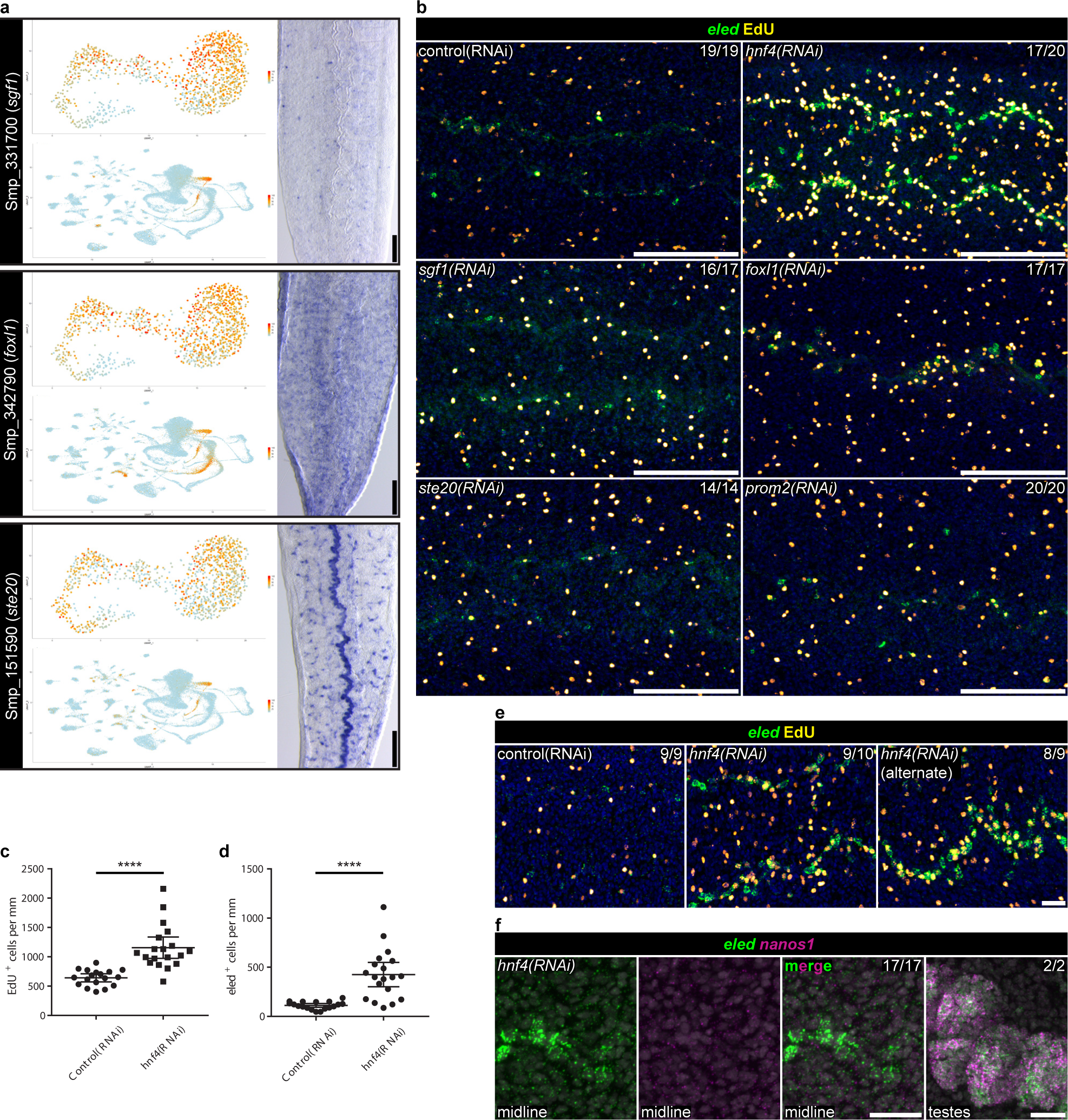
An RNAi screen identifies *hnf4* as a regulator of *eled*^+^ neoblasts **a**, For each of the “*eled*^+^ neoblast”-enriched genes *sgf1*, *foxl1*, and *ste20*: (top) UMAP projection plots of the expression pattern of the indicated gene on the re-clustered dataset from Fig. 3a, (bottom) on the entire dataset, and (right) a representative micrograph of colorimetric WISH of the indicated gene. **b**, Representative micrographs of FISH of *eled* in conjunction with an EdU pulse showing the location of *eled*^+^ neoblasts (green) and EdU^+^ proliferative cells (yellow) in the indicated RNAi condition. Target gene name is indicated in the upper left and the number of parasites similar to the representative micrograph is indicated in the upper right of each panel. Data are from two biological replicates. Nuclei are pseudo-colored blue. **c**-**d**, Graph showing quantification of the number of EdU^+^ proliferative cells (**c**) or *eled*^+^ cells (**d**) per mm of parasite from Fig. 3C in either control(RNAi) or *hnf4(RNAi)* animals. *n* = 18 control(RNAi) and 19 *hnf4(RNAi)* animals from two biological replicates. **e**, Representative micrographs of FISH of *eled* in conjunction with an EdU pulse showing the location of *eled*^+^ neoblasts (green) and EdU^+^ proliferative cells (yellow) in either control RNAi conditions (“control RNAi”), *hnf4* RNAi conditions (“*hnf4(RNAi)*”), or *hnf4* RNAi conditions using a separate, non-overlapping construct (“*hnf4(RNAi*) (alternate)”). The number of parasites similar to the representative micrograph is indicated in the upper-right of each panel. Data are from one biological replicate. Nuclei are pseudo-colored blue. **f**, Representative micrographs of double FISH of *eled* and *nanos1* demonstrating no co-expression along the parasite’s midline but strong co-expression of *eled* and *nanos1* in reproductive organs like the testes in *hnf4* RNAi conditions. The number of parasites grossly similar to the representative micrograph is indicated in the upper-right of each panel. *n* = 17 *hnf4(RNAi)* animals from two biological replicates. Nuclei are pseudo-colored grey. Scale bars, **a**, 100µm, **b**, 100µm, **e**, 20µm, **f**, 20µm. UMAP projection plots are colored by gene expression (blue = low, red = high). ****, *p*<0.0001 (Welch’s t-test).

**Extended Data Fig. 9.**
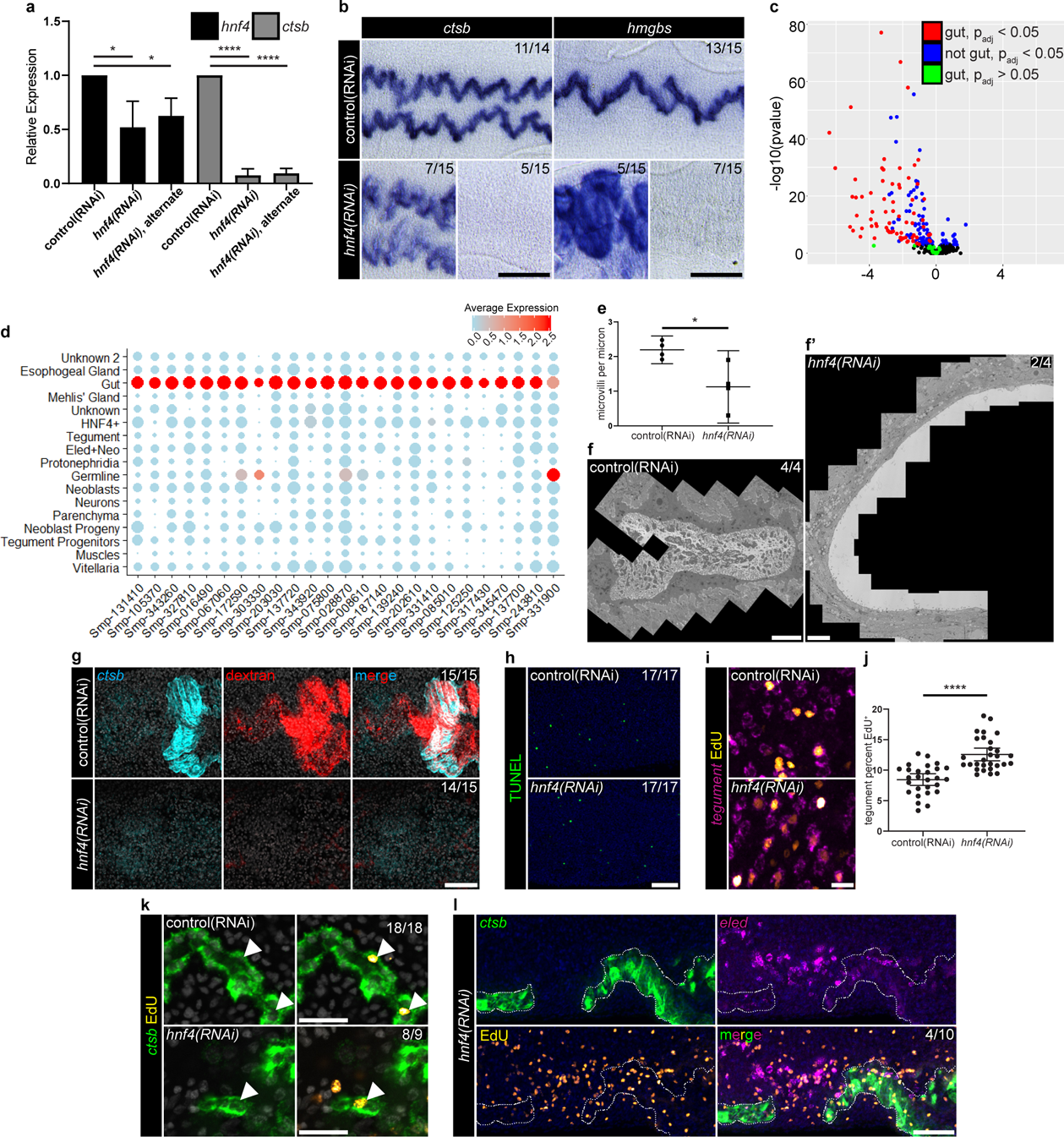
*hnf4* RNAi results in transcriptional and structural gut abnormalities **a**, Graph of relative quantification of *hnf4* mRNA (black) or *ctsb* mRNA (grey) as determined by qPCR in either “control(RNAi)”, “*hnf4(RNAi)”, or “hnf4(RNAi)* alternate*”* animals as. Data are from four biological replicates. **b**, For the “gut”-specific genes *ctsb* and *hmgbs*: Representative micrographs of colorimetric WISH of the indicated gene in either control RNAi conditions or *hnf4* RNAi conditions. The number of parasites grossly similar to the representative micrograph is indicated in the upper-right of each panel. Data are from three biological replicates. **c**, Volcano plot of data from an RNAseq experiment comparing gene expression of control(RNAi) animals to that of *hnf4(RNAi)* animals. “gut”, genes expressed in the “gut” cluster, “not gut”, genes not expressed in the “gut” cluster. **d**, A dot-plot summarizing the cluster-specific expression of each of the top 25 down-regulated genes in *hnf4(RNAi)* animals. Cluster IDs are on the vertical axis and gene IDs are on the horizontal axis. Expression levels are color by gene expression (blue = low, red = high). **e**, Graph showing quantification of the number of microvilli per micron of gut surface from Fig. 3e. Numbers are the average of 4 different sections of gut from each of 4 animals. **f**, Stitched TEM micrographs from either control(RNAi) animals **f,** or *hnf4(RNAi)* animals **f’**. The number of parasites similar to the representative micrograph is indicated in the upper-right of each panel. Data are from four animals from 2 biological replicates. **g**, Representative micrographs of double FISH of the gut marker *ctsb* (cyan) and the presence of fluorescently-labeled dextran (red) in the gut lumen in either control RNAi or *hnf4* RNAi conditions. The number of parasites grossly similar to the representative micrograph is indicated in the upper right of each panel. Data are from three biological replicates. Nuclei are pseudo-colored grey. **h**, Representative micrographs of a fluorescent TUNEL experiment showing apoptotic cells (green) in either control RNAi conditions or *hnf4* RNAi conditions. The number of parasites grossly similar to the representative micrograph is indicated in the upper right of each panel. Data are from two biological replicates. Nuclei are pseudo-colored blue. **i**, Representative micrographs of FISH for a pooled mix of four tegument-specific mRNAs^13^ (magenta) in conjunction with an EdU pulse followed by a 7 day chase showing the location of EdU^+^ progeny cells (yellow) in either control RNAi conditions or *hnf4* RNAi conditions. **j**, Graph showing quantification of the percentage of tegument cells that are EdU^+^ from **i**. *n* =27 control(RNAi) and 29 *hnf4(RNAi)* parasites from 3 biological replicates. **k**, Representative micrographs of FISH of the gut marker *ctsb* (green) in conjunction with an EdU pulse followed by a 7-day chase showing location of EdU^+^ progeny cells (yellow) in either control RNAi conditions or *hnf4* RNAi conditions. The number of parasites similar to the representative micrograph is indicated in the upper right of each panel. Data are from 2 biological replicates. Nuclei are pseudo-colored grey. **l**, Representative micrographs of double FISH of the gut marker *ctsb* and *eled* in conjunction with an EdU pulse showing the location of EdU^+^ proliferative cells (yellow) in *hnf4* RNAi conditions. The dashed line indicates the approximate boundary of the residual gut-like tissue found in *hnf4(RNAi)* animals. The number of parasites similar to the representative micrograph is indicated in the upper right of lower right panel. Data are from 2 biological replicates. Nuclei are pseudo-colored blue. Scale bars, **b**, 100µm, **f**, 5µm, **f’**, 5µm, **g**, 50µm, **h**, 50µm, **i**, 10µm, **k**, 20µm, **i**, 50µm. *, *p*<0.05 (Welch’s t-test), ****, *p*<0.0001 (Welch’s t-test).

**Extended Data Fig. 10.**
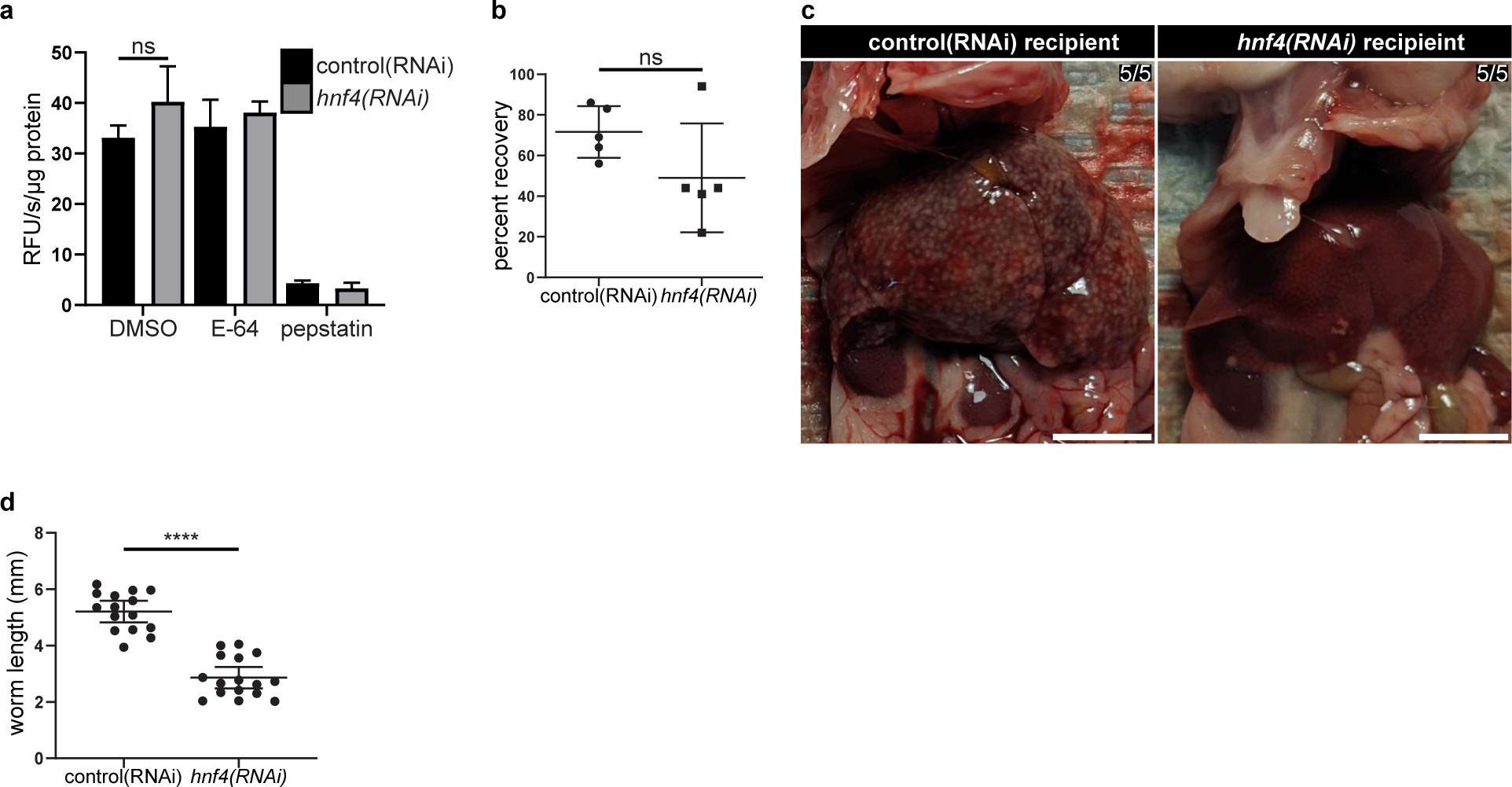
*hnf4* is required for blood feeding **a**, Graph of the aspartyl protease activity of lysates from control(RNAi) or *hnf4(RNAi)* parasites as determined by the ability to cleave the fluorogenic substrate, mca-GKPILFFRLK-K(dnp) in the presence of no inhibitor (DMSO), the general cysteine protease inhibitor E-64 (E-64), or the aspartyl protease inhibitor pepstatin A (pepstatin). **b**, Graph quantifying the recovery rate of worms from transplant recipients. Data are from five recipients. **c**, Representative photographs of livers of mice 30 days after transplant with RNAi-treated parasites. The number of livers grossly similar to the representative photograph is indicated in the upper right each panel. Data are from two recipients in one biological replicate. **d**, Graph showing quantification of worm length from Fig. 4f. *n* = 15 for control(RNAi) male parasites and 16 *hnf4(RNAi)* male parasites from 3 separate recipients. Scale bar, **c**, 1cm. ns, not significant, ****, *p*<0.0001 (Welch’s t-test).

